# A Unified Account of Lightness Illusions via Edge-Based Reconstruction of Natural Images

**DOI:** 10.64898/2026.04.07.716245

**Authors:** Srijani Saha, Talia Konkle, George A. Alvarez

## Abstract

The human visual system transforms patterns of light into rich perceptual experiences, where what we see is a construction that goes beyond simple measurement. Lightness illusions—where identical parts of an image can appear dramatically different depending on context—provide a window into these processes. Here we leverage a deep learning framework to investigate the constructive processes that give rise to lightness illusions, introducing the core computational goal of edge-based image reconstruction. Specifically, we demonstrate that autoencoder models trained to reconstruct natural images based only on an edge-based image representation naturally recapitulate a wide range of lightness illusions, which were previously assumed to require distinct mechanisms, inference over lighting sources, and explicit three-dimensional scene representation. These results offer a simpler, unified account of diverse lightness phenomena as emerging naturally from surface filling-in mechanisms, and broadly provide a framework for understanding the computational principles that underlie our perception of the visual world.

**SIGNIFICANCE STATEMENT:** The human visual system shows remarkably stable perception of objects under different viewing conditions, but it uses strategies that can be thwarted by clever visual illusions – for instance, the exact same object can appear as either white or black in different contexts. The most complex of these lightness illusions have long been taken as evidence that perception involves explicit inference about 3D scene geometry and lighting conditions. However, here we show that these illusions also emerge in deep learning models, trained simply to reconstruct natural images from sparse edge signals. Thus, our perception of the lightness of surfaces in our world may instead arise from a much more primitive computation — reconstructing surface appearance from edge responses.

## INTRODUCTION

When we look out at the world, our rich and detailed perceptual experience is delivered so fast and effortlessly that it gives the impression we somehow *directly perceive what is out there*. In fact, our perceptual experience is a mental construct, derived from local measurements of light on the retina, and processed through extensive neural circuitry of the visual system (Hubel & Wiesel, 1962; Kuffler, 1953; Sayim & Cavanagh, 2011). Indeed, the signals transmitted from the eye to the brain are nothing like the image entering the eye: e.g., retinal ganglion cells have center-surround receptive fields which respond primarily to local brightness contrast and color contrast (Kuffler, 1953; Rodieck, 1998). Apparently, the rest of the visual system must take these edge-based responses from the eye and transform them, ultimately constructing our rich perceptual experience of the scene geometry, the intrinsic lightness and colors of different surfaces and objects, and a sense of the ambient of illumination (e.g. high noon or golden hour) (see Figure 1A). How does the visual system reconstruct a detailed description of the surfaces of the world, starting from primarily local-contrast edge information?

**Fig. 1.**
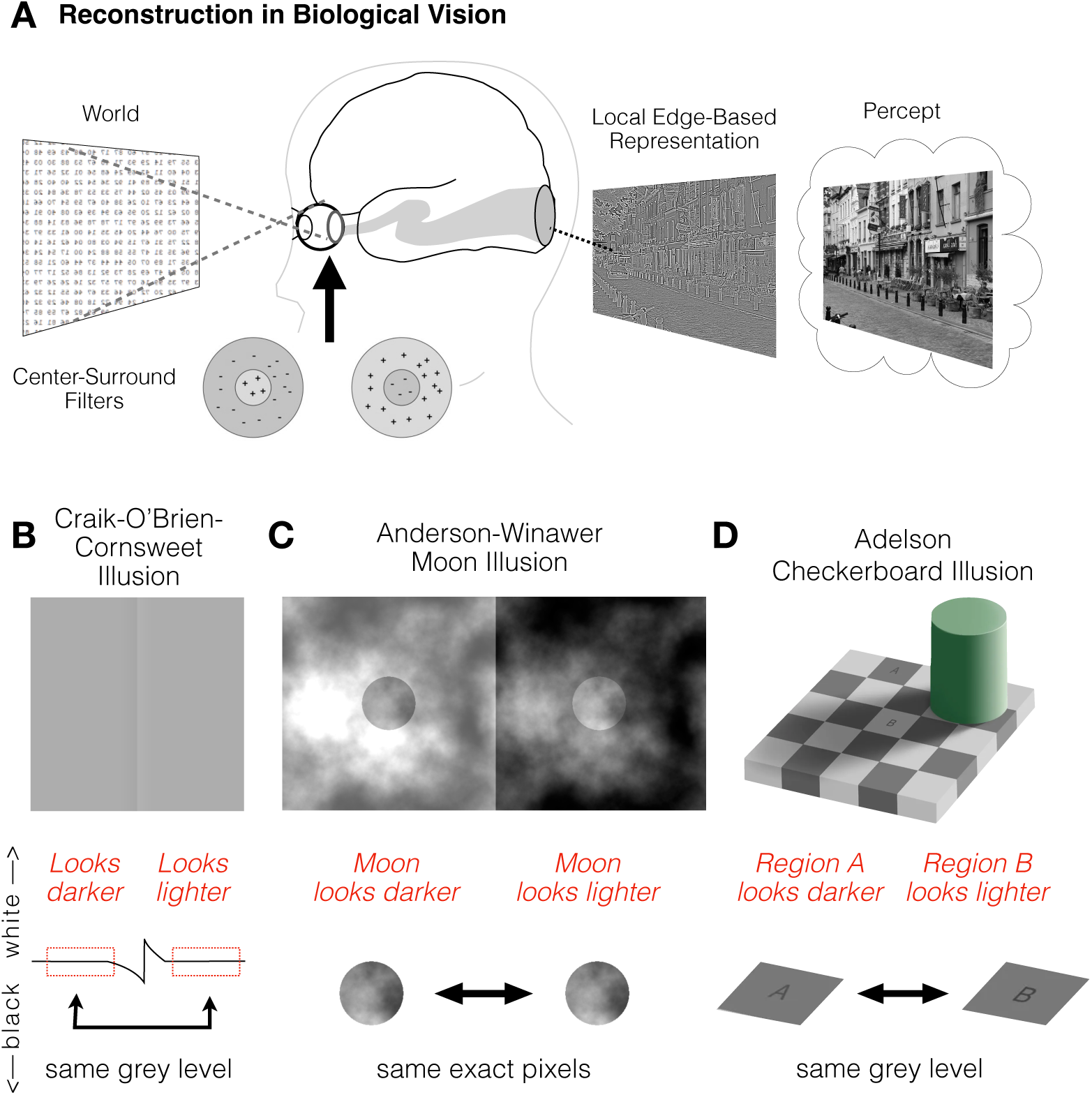
Reconstruction and lightness illusions in human vision. (A) In the biological visual system, center-surround filters as early in the retina transform measurements of light in the world into a contrast-edge representation. This means that what we perceive is not just the result of direct measurements from the world but rather a reconstruction from this edge-based input. The Craik-O’Brien-Cornsweet Illusion (B) the Anderson-Winawer Moon Illusion (C), and the Adelson Checkerboard Illusion (D) are the three primary illusions we will evaluate in Experiment 1. These illusions have been explained with mechanisms of increasing complexity, from edge-based mechanisms for the Craik-O’Brien-Cornsweet illusion, to the scene decomposition (object vs. ambient haze) in the moon illusion, to scene geometry and inferred lighting from shadows in the checkerboard illusion.

Lightness illusions have been a powerful tool for gaining insight into the reconstructive processes at work in the visual system. For example, consider the Craik-O’Brien-Cornsweet illusion (**Figure 1B**; Craik, 1966; O’Brien, 1958; Cornsweet, 1970), the Anderson-Winawer Moon Illusion **(Figure 1C**; Anderson & Winawer, 2005), and the Adelson Checkerboard Illusion (**Figure 1D**; Adelson, 1995). Henceforth, we refer to these illusions using a shorthand of Cornsweet Illusion, Moon Illusion, and Checkerboard Illusion. In the Cornsweet Illusion, we see a dark panel next to a light panel, but in fact both panels are the same medium gray; the surface brightness is being ‘filled in’ by our own perceptual systems, induced by a very thin light-dark contrast edge in the middle of the image. In the Moon Illusion, the moons appear either nearly white or nearly black, but in fact the pixels in the center disc-region of both images are exactly the same! Here, the different brightness levels of surrounding background and haze induce us to see the moon as either light or dark. In the Checkerboard Illusion, region A appears to be darker than region B, whereas in fact they are identical medium gray. Here, your visual system appears to take the 3D scene geometry and lighting conditions into account, inferring that region B is within a cast shadow yet reflects the same amount of light as region A in direct light, which could only occur if region B was a lighter surface. These illusions reveal that our perception is not a veridical reflection of directly measured light but instead is a highly constructive process that appears to depend on local contrast, context, 3D scene geometry, inferred lighting conditions, and prior knowledge in ways that are still not fully understood (Gregory, 1997; Brainard & Hurlbert, 2015; Gilchrist, 2015; Lafer-Sousa et al., 2015).

The Cornsweet Illusion must result from the visual system detecting the small luminance difference at the edge and effectively propagating that contrast difference throughout the adjacent surfaces (Grossberg & Todorović, 1988). However, it has been argued that more complex phenomena, like the Moon Illusion, cannot be explained by such simple filling-in mechanisms (Anderson & Winawer, 2005; Cornsweet, 1970), and that much more sophisticated scene decomposition and analysis are required to explain the Checkerboard Illusion (Adelson, 1995, 2000). An influential framework for understanding these more complex phenomena is provided by the “inverse graphics” theory of vision (Adelson, 1993; Barrow & Tenenbaum, 1978; Kersten, 1997). On this account, our perception of the world is based on inferential processes, where the goal of the visual system is to estimate the intrinsic properties of objects by factoring out ambient lighting conditions, and involves sophisticated mechanisms that take 3D scene geometry and assumptions about lighting sources into account (Gilchrist, 1977; Purves et al., 1999). For example, in the Moon Illusion, it is assumed that our visual systems must infer a hazy surface in front of the moon and then factor out the haze to estimate the moon’s brightness, and in the Checkerboard Illusion it is assumed that 3D geometry and inferences about light sources and shadows are needed to explain the percept.

One significant challenge for inverse graphics accounts is that reverse engineering the world state and lighting sources is a classically ill-posed problem, and doing Bayesian inference over all possible scene structures and lighting sources to infer the most likely world state is computationally intensive even with proposed solutions to reduce the search space (Yuille & Kersten, 2006) —there is currently no computational model of this kind that we know of that operates over natural images and can explain lightness illusions. Instead, there has been advances inspired by inverse graphics framework — such as Bayesian Intrinsic Image Models (Allred & Brainard, 2013; Murray, 2013, 2020) and Equally Illuminated Models (Bloj et al., 2004; Brainard & Maloney, 2011)— which can be fit to behavioral judgements of simple lightness illusions or scenes; however, these models do not operate over complex, natural images.

In the current work, we consider an alternative hypothesis that a local edge-based filling-in mechanism is sufficient to account for how we subjectively ‘paint’ our world and reconstruct intact surfaces from edge responses. On this account, local light and dark measurements at the contrast edges alone may be sufficient to account for a diverse set of lightness illusions, without requiring an explicit inferred model of the full scene geometry and lighting sources (Blakeslee & McCourt, 1999; Cohen-Duwek & Spitzer, 2018, 2019; Dakin & Bex, 2003; Rudd, 2014; Cornsweet, 1970). While there have been some efforts to explore these ideas computationally in the past (Land & McCann, 1971; Blakeslee & McCourt, 1999; Dakin & Bex, 2003), here we take advantage of recent breakthroughs in deep learning and computer vision, leveraging models whose computational goal is to directly to reconstruct images (Goodfellow et al., 2014; Hinton & Salakhutdinov, 2006; Ho et al., 2020; Kingma & Welling, 2022; Ronneberger et al., 2015; Vincent et al., 2008). Specifically, we passed images through a Laplacian edge filter, providing pseudo “retinal ganglion cell responses” (Marr & Hildreth, 1980; Rodieck, 1965), and then trained an autoencoder (Ronneberger et al., 2015; Wang et al., 2020) to reconstruct the full-intact original image from these edge responses (**Figure 2A and 2B**). Thus, the computational goal of this model system is to “paint-in” the rest of the pixel information based on local-contrast edge-responses alone. We then take this model, trained only on natural images, and probe it with images that induce lightness illusions in people, and measure whether the model’s reconstruction errors echo perceptual illusions. Critically, we do not consider this to be a biologically plausible model — instead we consider it to be an abstraction of the task achieved by human vision (reconstructing surface lightness given edge-responses from the retina). In this way, we leverage the model as a testbed for understanding what kinds of lightness phenomena can occur purely from the simple goal of edge-based image reconstruction.

**Fig. 2.**
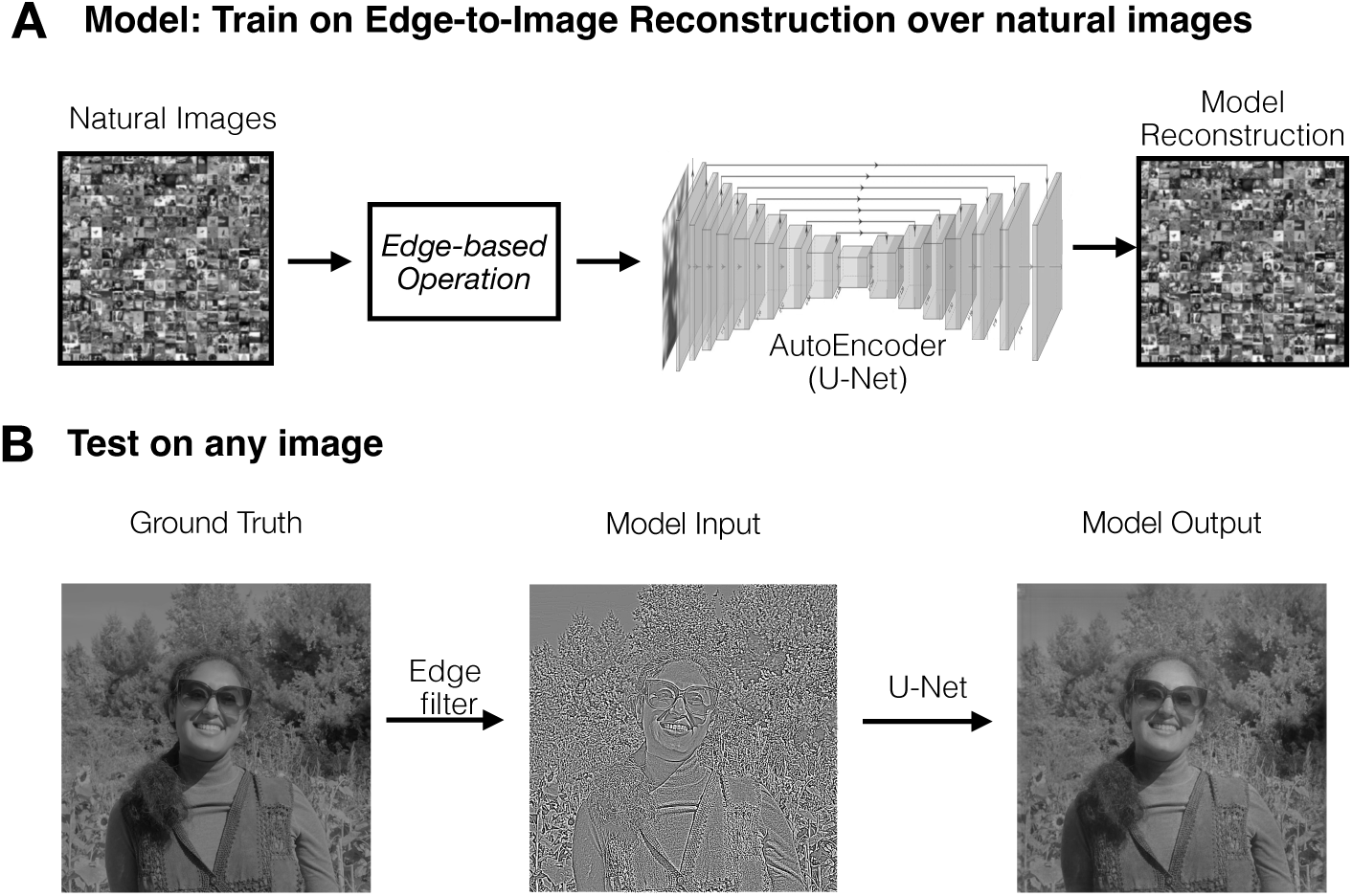
Edge-based Reconstruction in a deep neural network model. (A) We used a U-Net model to operationalize this high-level computational goal, training the model to take in edge-based inputs of natural images and to reconstruct the full gray-scale images. (B). Example reconstruction of a new (untrained) image: the original image is shown, followed by the result of the edge filter, which was the input to the model, along with the model output image. This image is a self-portrait of the corresponding author, used with permission.

Here, we first demonstrate that these “Edge-Net” models can in fact succeed at accurate edge-based image reconstruction of natural images. We then present these models with a variety of lightness illusions (also passed through an edge filter), to explore how the model reconstructs these images, based on what it has learned from reconstructing natural images. Note the models were never trained on illusions; we are probing for purely emergent phenomena due to learning to reconstruct natural images from edge-based input. To anticipate the results, we find that models trained only on reconstructing natural images from edges can naturally recapitulate all three illusions (Cornsweet, Moon, Checkerboard). We additionally show that simple denoising reconstruction goals do not show the same emergent illusions. We then explore a broader suite of illusions where the edge-based reconstruction model succeeds and identify boundary conditions where the model does not mimic human perceptual illusions. Finally, we provide a direct quantitative comparison between humans and models on the Moon illusion and show that the model aligns both qualitatively and quantitatively with human perceptual judgments. Based on these results, we argue that lightness illusions need not reflect an explicit goal of uncovering intrinsic properties of objects; instead, we offer that edge-based reconstruction is a modular goal (encapsulated to early visual processing), and that this objective alone results in outputs that are biased towards depicting intrinsic object properties that remain constant across lighting conditions.

## RESULTS

### Edge-Based Reconstruction Model Successfully Reconstructs Natural Images

We introduce an “Edge-Net model” as is a U-Net model trained to reconstruct natural images from inputs containing local-contrast edge-responses only. The architecture was a standard U-Net taken from Wang et al. (2020) and consisted of an 8-layer encoder, 2048-dimensional bottleneck, and an 8-layer decoder. To train the model, we first sampled a random set of 1024 images from the ImageNet dataset across the 1000 categories and processed them with a standard edge-based filter of a second-order gradient (i.e., a Laplacian filter). This Laplacian filter acts as a band-pass filter which approximates the output of a center-surround receptive fields of the retina and LGN (Marr, 2010). In this way the output image is transformed into a map of “contrast edges” that have a lighter and darker side; note this is not the same as a line drawing representation of binary edges (e.g. as you might get from a Canny filter edge detector). These edge-response inputs were passed into the model, where the objective for the model is to output a grayscale image and minimize the pixel difference between the output images and the original images (see **Figure 2A**). This was accomplished through a loss function following the procedure of Wang et al. (2020), which computed the mean absolute error between the original grayscale image and the reconstruction, over the final output stage and all intermediate outputs of the decoder blocks (see Methods).

The Edge-Net model was successfully able to reconstruct images from these edge-based inputs. Qualitatively, Figure 2B highlights the model’s reconstruction for an image not present in the ImageNet dataset – a self-portrait– as testament to the model’s reconstruction capacity for natural images more broadly. Quantitatively, the model achieved a high reconstruction accuracy on the held-out validation set of 300 images, with an average mean absolute error of only 5.4% ± 0.13% (standard error; see also Supplementary Fig. 1). Given this successful edge-based reconstruction, this model serves as an interesting proxy for the computational problem faced by the biological system to go from LGN-like edge responses to a filled-in percept.

### Edge-Based Models can reproduce the Cornsweet, Moon, and Checkerboard Illusions

Next, we examined whether the process of reconstructing images from edge-responses would result in systematic errors that mirror human lightness illusions. Would the model “paint” regions darker when humans perceive them as darker, and paint them lighter when humans perceive them as lighter?

To assess the robustness of any alignment between model reconstruction errors and human perceptual illusions, we trained a total of 32 Edge-Net models across four training datasets, varying the random initialization seed for each run (eight seeds per dataset) with comparable reconstruction performance within each set (see Supplementary Fig. 2). We specifically tested variations in the natural-image diet used to train the U-Net by sourcing two smaller datasets (1k_ds1, 1k_ds2; 1024 images each) and two larger datasets (8k_ds1, 8k_ds2; 8,000 images each), with minimal overlap between one of the smaller and larger datasets (see Methods for details).

First, we tested whether these edge-based reconstructive models can reproduce the illusory percept in the Cornsweet Illusion. The illusion image is passed through the edge-based filter into each of the trained model (**Figure 3A**). If a model paints in the region we perceive to be lighter (illusory light region) with a greater mean pixel value than the region we perceive to be darker (illusory dark region), then we have evidence that the model’s reconstruction errors reproduce the direction of the illusory percept. Accordingly, the key outcome measure of illusion magnitude is calculated as the difference between the mean pixel intensity in the reconstructed illusory light and dark regions, relative to a zero pixel intensity difference in the ground truth image. As shown in **Figure 3A**, all Edge-Net models consistently reproduced the illusion, across training datasets (1k_ds1: 8.9±1.4, t(7)=6.3, p<0.001; 1k_ds2: 11.6±1.1, t(7)=10.9, p<0.001; 8k_ds1: 5.1±0.8, t(7)=6.5, p<0.001; 8k_ds2: 5.0±0.7, t(7) = 7.4, p<0.001, one sample t-tests).

**Fig. 3.**
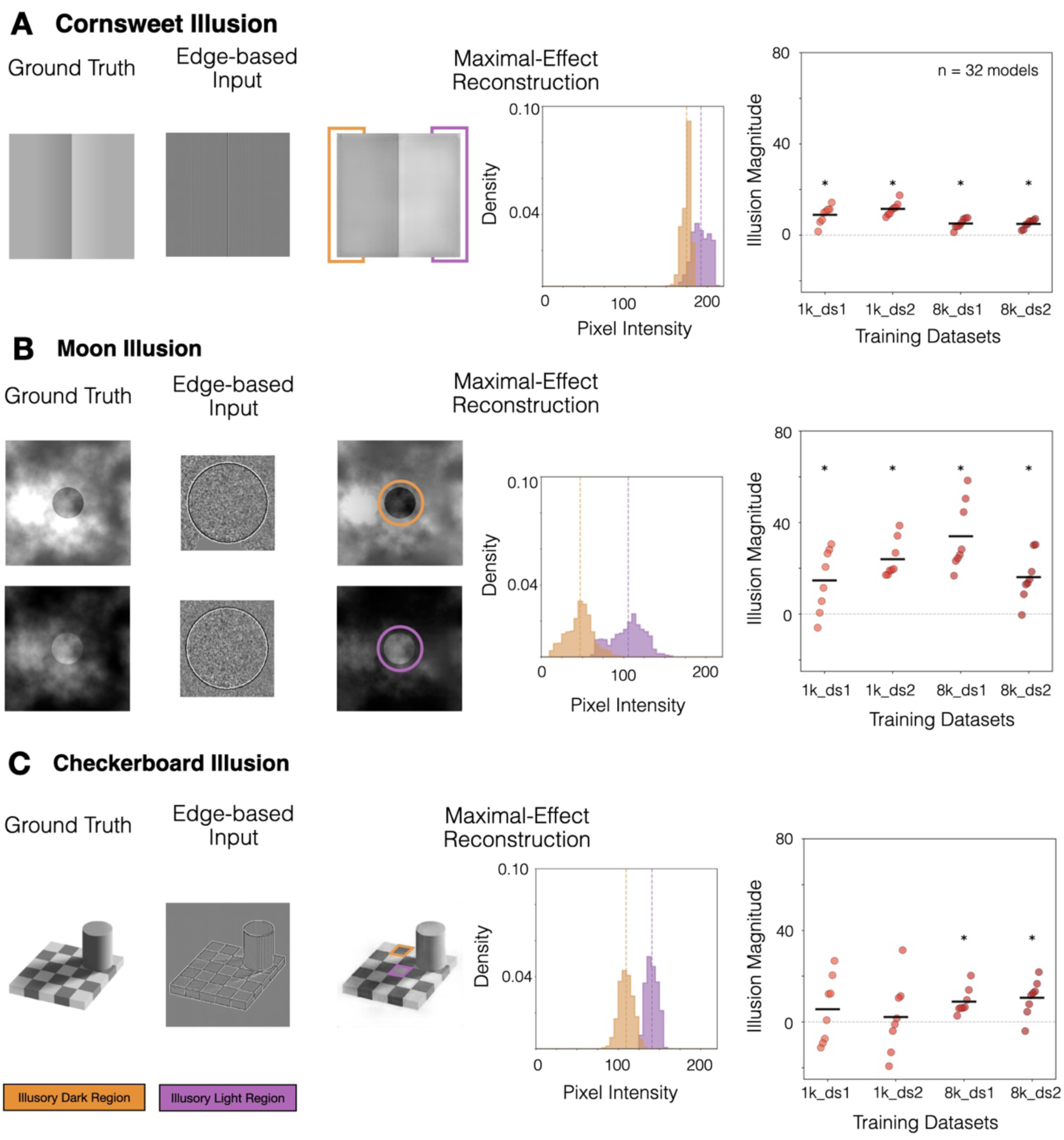
Edge-based Model Reconstructions of Cornsweet, Moon, and Checkerboard Illusions. For each illusion, we show the ground truth image(s), the contrast-edge image(s) after the Laplacian filter, and the model’s reconstruction for the model instance that showed the maximal illusion magnitude. The orange and purple annotations highlight the illusion’s “regions of interest”—they are equivalent intensity in the ground truth image but differ in human perceptual judgements. The adjacent plot shows the distribution of the pixel intensity in the regions of interest in the provided reconstruction on the standard 0-255 scale (x-axis). Finally, the last plot for each illusion provides the summary of the illusion magnitude performance (y-axis) across each subset of models by training dataset (x-axis). Each dot represents the illusion magnitude for each model, and each black bar denotes a dataset mean. For the Cornsweet and Moon Illusions, all sets of models reliably painted the illusory dark region as darker than the illusory light region. Models trained on the larger datasets more robustly reproduced the Checkerboard Illusion.

We next tested the Moon Illusion. Both illusory images – the dark moon in light haze and light moon in dark haze – were passed through an edge-based filter and reconstructed by each model (Fig. 3B). Here, the illusion magnitude is the difference in mean pixel intensity in the reconstructed moons. Although the moons are the exact same pixels in the two original images, Edge-Net models again reproduce the human perceptual illusion, across training datasets (1k_ds1: 14.7±4.9, t(7)=3.0, p=0.019; 1k_ds2: 23.4±3.0, t(7)=8.1, p<0.001; 8k_ds1: 34.0±5.3, t(7)=6.4, p<0.001; 8k_ds2: 16.1±3.7,t(7)=4.4, p=0.003). We did observe that the reconstructions for the Moon Illusion are noticeably darker than the original images. Additional analysis on quasi-natural images (additional set of hazes) and natural images suggest that the models exhibit a bias for darker reconstructions when edge strength is weak (see Supplementary Fig. 3). Regardless of the cause for this overall main effect, we focus here on the relative reconstruction difference within the key region of interest for the human illusory percept. A further analysis on a larger programmatic set of 500 variations of the Moon Illusion (see Methods and Supplementary Figure 4 for details) revealed that the Edge-Net models reproduce the illusion across a majority of haze structures (1k_ds1: 476/500, 1k_ds2: 389/500, 8k_ds1: 423/500, 8k_ds2: 380/500, all p < 0.05, one sample t-tests).

Finally, we tested whether the Edge-Net models could also reproduce the more structurally complex Checkerboard Illusion. After passing the image through the edge-based filter and then reconstructing it with each model, we calculated the illusion magnitude as the difference in the mean pixel intensity in the reconstructed illusory squares (Fig. 3C). We found that only the Edge-Nets trained on the larger natural image datasets systematically recapitulated the Checkerboard Illusion (8k_ds1: 8.9±2.0, t(7)=4.4, p=0.003; 8k_ds2: 10.5±2.8, t(7)=3.8, p = 0.007). The models trained on the smaller datasets did not (all p values > 0.05). Given that this illusion is more complex in scene structure than the Cornsweet and Moon Illusions, these results reveal that greater (though still modest) exposure to natural image statistics is required for the Checkerboard Illusion to emerge in the model’s reconstruction.

Broadly these results show that these Edge-Net models—trained solely to do edge-based image reconstruction of natural images—learn filling-in mechanisms that naturally recapitulate this range of lightness illusions. These results have powerful implications, as there was no specific training tailored to each of these illusions; instead, errors mimicking these illusions naturally emerged from a general learning objective operating over natural image input. Further, while traditionally it has been assumed that the more complex Moon Illusion and Checkerboard Illusion would require more sophisticated scene processing and inference mechanisms than the simpler Cornsweet Illusion, here we show that models trained with one objective (edge-based reconstruction) naturally mimic the illusory percept for all three illusions using domain-general mechanisms.

### Denoising reconstruction does not elicit lightness illusions

Is it the case that any reconstructive or generative goal will result in errors that mimic human lightness illusions? To address this question, we tested whether denoising models which reconstruct clean images from noisy inputs, also mimic these illusions. This comparison is theoretically meaningful because both edge-based reconstruction and denoising involve learning to ‘fill in’ missing information based on natural image statistics (Hyvärinen et al., 2009). However, they differ fundamentally in what information is preserved: edge filters maintain local contrast relationships while discarding absolute intensities, whereas noise corruption preserves degraded intensity information while disrupting fine spatial details. If lightness illusions emerge from any reconstruction process that learns natural image priors, denoising models should also exhibit them. Alternatively, if these illusions specifically emerge from mechanisms that support reconstructing from contrast edges — as suggested by neurophysiological evidence (Hung et al., 2007) and psychophysical evidence (Rudd, 2014; Kingdom, 2011) — then denoising models should not recapitulate these effects despite their similar computational goal of reconstruction.

As shown in Fig. 4A, we trained a set of “DeNoise-Nets”, using the same U-Net architecture as in the previous experiments, with 8 models trained to remove low levels of gaussian noise (std = 0.10 for intensity values on -1 to 1 scale) and 8 models to denoise high levels of gaussian noise (std = 0.30). To enable a controlled comparison between reconstructive objectives, we trained the DeNoise-Nets on the same dataset (8k_ds1) for which Edge-Net models showed robust and consistent effects across the three illusions (see Methods for further training details). Based on mean absolute error, reconstruction quality was comparable between denoising and edge-based reconstruction models, with modest differences in dynamic range (Supplementary Fig. 5), indicating the trained DeNoise-Nets can effectively remove noise.

**Fig. 4.**
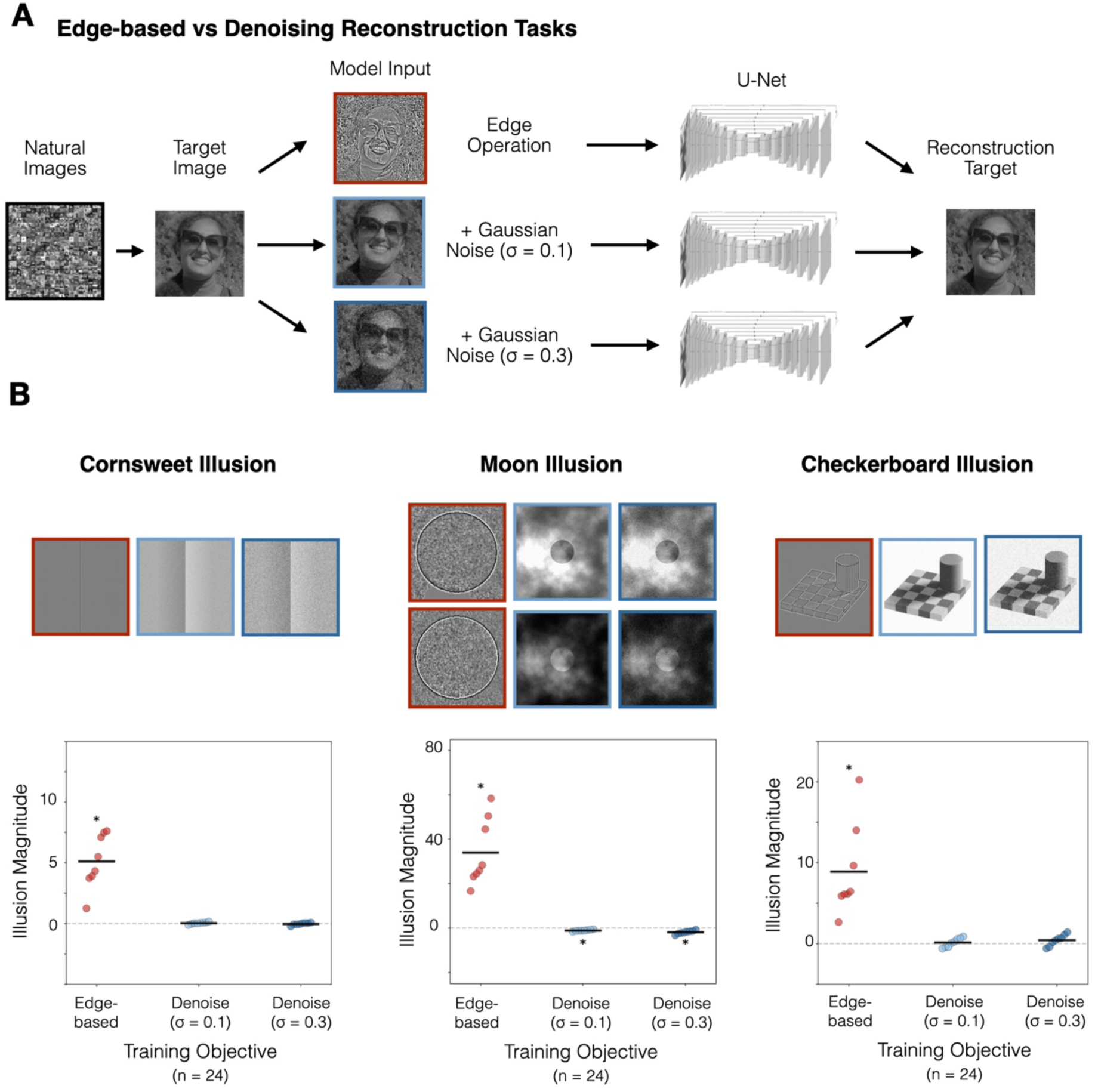
Comparison of the Reconstructions of the Cornsweet, Moon, and Checkerboard Illusions between the Gaussian Denoising Models and the Edge-Based Reconstructive Models. (A). Training pipeline for matched U-net models, trained on natural images that are either processed into contrast-edges or are corrupted by low or high levels of pixel-wise Gaussian noise. The reconstruction goal of all models is to recover the original grayscale images. Models share the same input diet (8k_ds1), architecture, and output target, and vary only in terms of the input processing. Example input image is a self-portrait of the corresponding author, used with permission. (B). Average reconstruction results for the Cornsweet Illusion, Moon Illusion, and Checkerboard Illusion. For each illusion, the inputs across each model type are presented above a summary plot. The three model types (edge-based, low-noise denoising, and high-noise denoising) are shown along the x-axis of the summary plot. Each dot represents the illusion magnitude for each model, and each black bar denotes a dataset mean. The set of edge-based models reliably show a stronger effect of each illusion in relation to either set of denoising models.

Figure 4B shows the DeNoise-Net models’ reconstruction performance on the three illusions, in the context of the Edge-Net models trained on the same dataset. To evaluate the illusion strength in the denoising models, each illusion was passed through the denoising models ten times with different noise patterns, and we evaluated the models’ performance on the average reconstruction. Overall, the denoising models did not show emergent illusory reconstructions. For the simple Cornsweet Illusion, the models produced veridical reconstructions (mean pixel intensity difference: low-noise: 0.04±0.02, t(7)=1.2, p>0.05; high-noise: -0.03±0.03, t(7)=-0.8, p>0.05). For the Moon Illusion, the denoising models actually showed a very small but systematic *reversal* of the expected illusion direction, reconstructing the illusory darker moon as slightly brighter on average (low-noise: -1.1 ±0.1, t(7)=-9.3, p<0.001; high-noise: -1.9±0.3, t(7)=-6.7, p<0.001). And, for the Checkboard Illusion, the denoising models again showed veridical reconstruction (low-noise: 0.1±0.2, t(7)=0.7, p>0.05; high-noise: 0.4±0.2, t(7)=1.8, p>0.05). We additionally trained a set of low-noise and high-noise denoising models on the smaller dataset of 1k_ds1 and only observed at most weak and inconsistent illusion effects across all three cases, with substantially smaller illusion magnitudes than those produced by edge-based models (Supplementary Fig. 6).

These denoising model results show that not just any reconstructive goal is sufficient for a system to show these emergent lightness illusions. These results imply that, for these illusions, explicit contrast-edge information plays a critical role in providing the computational constraints that can lead the model to paint surfaces with systematic errors that mirror human lightness illusions.

### Edge-Based Reconstruction Across a Diverse Set of Lightness Illusions Reveals Systematic Generalization and Limits

To establish boundary conditions for which illusions are recapitulated by the edge-based reconstruction models, we next tested a broad suite of lightness illusions shown in Figure 5 (see Supplementary Fig. 7 for the full set of quantitative results).

**Fig. 5.**
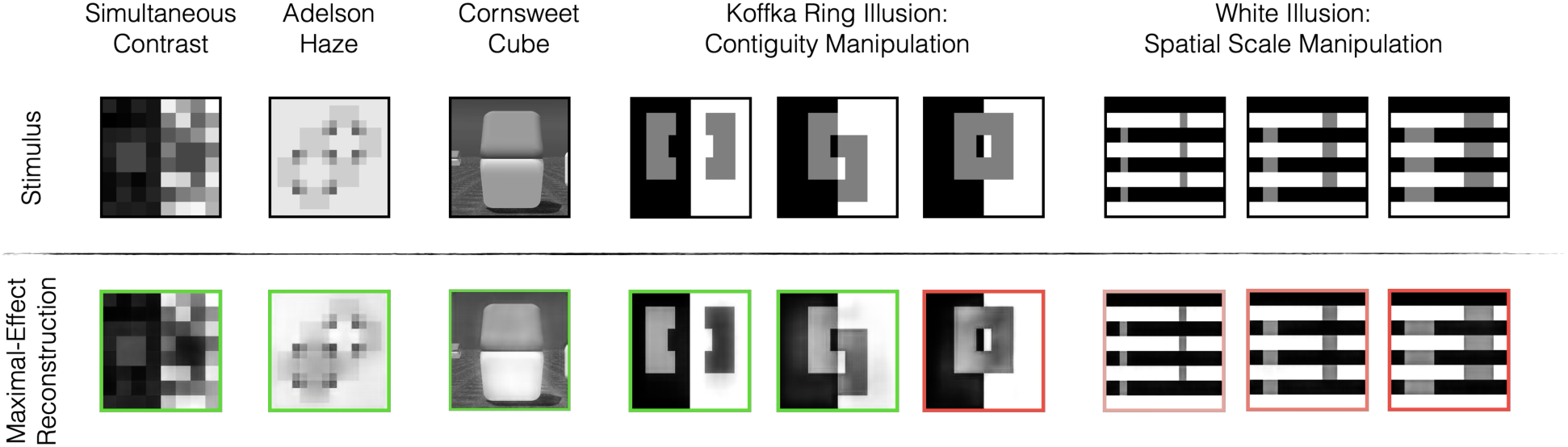
Edge-based Reconstruction Performance Across an Extended Suite of Illusions. Several classic brightness illusions are recapitulated by the model (reconstructions with the green border show errors in line with human percepts), including simultaneous contrast, the Adelson Haze illusion, the Cornsweet Cube Illusion, and some variants of the Koffka Ring and White Illusion. Reconstructions that do not align with human percepts (red border) include the Koffka Ring when the “[” and “]” regions are connected, and the White Illusion (see Footnote 1).

In general, we find that a diverse set of lightness illusions are recapitulated by Edge-Net models, including the Simultaneous Contrast Illusion, the Adelson Haze Illusion, the Cornsweet Cube Stimulus and flattened variants (including sensitivity to edge orientation). Each of these illusions has led to fairly distinct and tailored hypotheses about the computations of the visual system. For example, the Adelson Haze Illusion has been interpreted as evidence that the visual system must estimate and invert ‘atmospheric transfer functions’—complex mappings between surface reflectance and luminance that account for viewing conditions like haze or transparency (Adelson, 2000; Somers & Adelson, 1997). The Cornsweet variants, particularly sensitivity to edge orientation, have been used to argue that the human visual system infers 3D structure and assumes lighting from above in its construction of surface lightness. However, our general Edge-Based reconstruction models, trained on natural images, naturally make errors on these illusory stimuli that mimic human illusory percepts. And, the DeNoise-net models largely failed to recapitulate any of these illusions, apart from weak effects for the Adelson Haze Illusion and some flattened Cornsweet variants (see Supplementary Fig. 7). Together, these results indicate that reconstructing from contrast edges is sufficient to capture substantial structure in human lightness perception, and that many perceptual phenomena may emerge from this simpler, more local computational mechanism.

We also identified several telling mismatches between edge-model reconstructions and human illusory percepts, which suggest a systematic limitation. Specifically, if a region to be filled-in is surrounded by both equally and opposite dark and light edge influences, the models do not appear to resolve this competition the same as the human perceptual system. For example, in the Koffka illusion, the models broadly reconstructed the two disconnected C-shaped blocks as light and dark, in line with human perception; but when the two C-shape blocks were connected, the models still reconstructed the left lighter and the right darker, whereas humans perceive the central ring as uniform gray. A similar failure mode is evident in the parametric variations of the White Illusion. People see the left vertical stripe as lighter than the right vertical stripe for all “widths.” Models, on the other hand, never consistently show the illusion and in fact progressively show an anti-illusory pattern as the width increases^1^. These cases suggest a systematic limitation of these models, reflecting conditions where local edge information, in this implementation, is insufficient to resolve competing perceptual cues exactly as humans do. We discuss this further in the General Discussion.

### Human Behavioral Signatures of the Moon Illusion

In the previous sections, we focused on categorical comparisons between well-known, and highly consistent perceptual illusions, and the reconstruction errors of edge-based (and denoising) reconstructive models. One reason for this approach is that it is difficult to match the “task” performed by people in typical lightness experiments with the model’s reconstruction task. Here, we develop a new method that enables a more direct comparison between humans and models, focusing on the moon illusion.

To provide a direct comparison between “model reconstructions” and human judgments, we presented humans with target illusory stimuli and had them perform their own “reconstruction” by adjusting the appearance of an adjacent moon display in order to match their percept (see Fig. 6A and Methods for more details). Participants moved the mouse in a two-dimensional response space to adjust the overall brightness and contrast of pixels within the moon region (“Pattern Match Task”). The true values of the overall brightness and standard deviation of brightness of the target moon was one of the choices in the 2-D response space, so a veridical report of the pixel pattern is possible. Now, like the edge-based reconstructive models, participants have the same objective – to adjust the pixels within the moon to match the target – allowing us to test whether humans and models show comparable error magnitudes. For controlled comparison to prior work (Anderson & Winawer, 2005), we also conducted a different variant of the task where the moon is a uniform color (no haze), and participants slide one-dimensionally from black-to-white to report the moon lightness (“Uniform Match Task”).

**Fig. 6.**
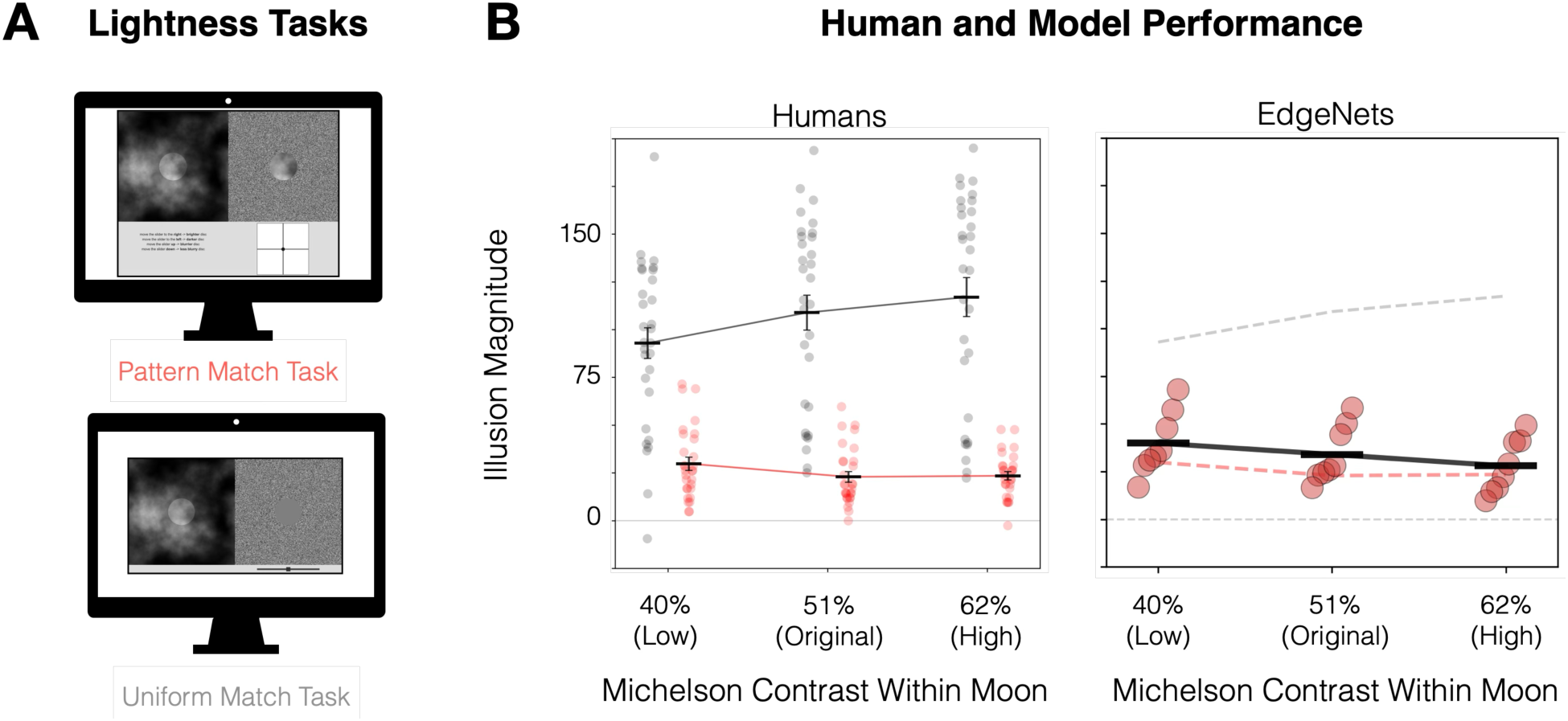
Human and Model Signatures of Anderson-Winawer Moon Illusion and the parametric effects of contrast. A) In the human behavioral “Uniform Match Task” and the “Pattern Match Task,” participants were shown the target stimulus on one side of the screen and had to either match their percept of the moon to a uniform moon or a perceived patten of the moon on the other side of the screen respectively. B) The left panel shows the illusion magnitude (y-axis) for participants as a function of the Michelson Contrast of the moon (x-axis), and each dot is a single human participant. The right panel shows a comparison between average human behavior (uniform match: gray dashed line; pattern match: red dashed line) versus 8 edge-based models (red dots). The illusion magnitude (y-axis) is again plotted as a function of the Michelson Contrast (x-axis) of the moon in the target image for the same stimuli. Each dot represents a model instance, and each black bar represents the mean performance across models.

Figure 6B plots the illusion magnitude reported by a group of 29 human participants in the Pattern Match Task (gray dashed line), 30 human participants in the Uniform Match task (red dashed line), and 8 edge-based models trained on 8k_ds1 (one red dot per model), as a function of contrast within the moon region (see Supplementary Fig. 8 for visualizations of participant and model responses). Replicating prior work (Anderson & Winawer, 2005), in the uniform match task, illusion strength in people *increases* with contrast. However, when people perform the pattern match task, we observed that illusion strength *decreases* with contrast. The Edge-Net models paralleled human illusion strength in the pattern match task, showing comparable illusions strength and similar reduction in illusion strength with increasing contrast. The same pattern of results was observed across different training datasets for the models (see Supplementary Fig. 9), indicating that when the human task is more closely aligned with the model task, there is a strong correspondence between human and model performance.

These results also highlight the importance of how the effects are measured, since humans show quantitatively and qualitatively different trends in the uniform and pattern matching tasks. When the task is a more direct measure of reconstructing what you see (pattern matching), we find comparable illusion magnitude strength and sensitivity to contrast in the Edge-Net and people. This implies that the filling-in mechanism of Edge-Nets are sensitive to the strength of contrast edges in qualitatively similar ways as the human perceptual system.

## DISCUSSION

The present study focuses on the fundamental question of why we perceive the world the way we do: Is it because our visual system is guided by a high-level computational objective, aiming to disentangle external light sources from the intrinsic properties of the surfaces around us (à la inverse graphics)? Or is it because our visual system is tasked with a more primitive, modular goal, aiming to construct a complete percept from an efficient and compressed, edge-based code? To address this question, we trained the Edge-Net model, a U-Net architecture optimized over natural images to reconstruct grayscale image from edge-based responses, and then evaluated the model on test images to see whether the model makes systematic errors that mimic human lightness illusions. Despite never encountering the illusion stimuli during the training, edge-based reconstructive models broadly recapitulated a wide-range of lightness illusions. Moreover, edge-based reconstructive models paralleled human sensitivity to the Moon Illusion in direct quantitative comparisons. Critically, models trained on the same datasets to remove Gaussian noise — with identical architecture and training images — did not consistently recapitulate the lightness illusions.

These results suggest that a single, modular learning goal—fill in luminance from contrast edges under natural-image priors—can account for a substantial subset of classic lightness phenomena. Conceptually, lightness illusions here appear as reconstruction by-products of an “edges-in → surfaces-out” objective rather than explicit optimization targets. Below we connect these results to classic accounts of lightness illusions and propose future work that could strengthen the alignment between Edge-Net reconstructions and human percepts.

### Relevance to theories of lightness perception

Our work starts with the premise that the vast majority of information transmitted from the retina to the cortex comes from neurons that respond only to local discontinuities in brightness and color, i.e., local edge responses (Kuffler, 1953; Rodieck, 1998), a form of compression that is perhaps necessary due to the information bottleneck of the optic nerve (Lindsey et al., 2019). Given that we ultimately perceive complete, intact surfaces, the human visual system appears to have mechanisms for reconstructing surface representations from these edge responses. Many prior works have focused on this puzzle, most notably the canonical “filling-in” multiscale spatial filtering model of Grossberg and Todorović (1988), which provides a detailed mechanistic account of how boundary-constrained lateral filling-in mechanisms can give rise to a wide array of brightness phenomena. While our work starts from a shared premise (filling-in is required), we do not provide a mechanistic account of the representations or network interactions necessary to achieve this goal. Instead, we leverage a deep-learning framework, in which we can specify the computational goal (desired input → output mapping), architectural constraints (multiscale U-Net), and visual diet (natural images), and allow the network to learn mechanisms that solve the task. This framework enables us to investigate lightness illusions not as the target of the model, which was trained only on natural images, but instead as a probe to see what behaviors naturally emerge as a by-product of the true goal of the system — to accurately reconstruct surfaces from edge responses in natural images. In short, Edge-Net operationalizes the “edges-in, surfaces-out” hypothesis at scale and lets us ask which illusions follow from that objective alone.

While an edge-based filling-in mechanism seems capable — possibly necessary — to explain the Cornsweet illusion, more complex lightness illusions have typically been assumed to require more explicit recovery of scene structure and ambient lighting, motivating inverse graphics theories of perception (Adelson, 1993; Barrow & Tenenbaum, 1978; Yuille & Kersten, 2006). Over the years, there has been extensive work along these lines, spanning from classic work like the Retinex model (Land and McCann, 1971), to more recent approaches like the Markov illuminance and reflectance “MIR” model (Murray, 2020). This inverse graphics framework differs from ours chiefly in the proposed computational goal: inverse graphics theories assume that the purpose/goal/design of the system is to discover and represent the intrinsic properties of objects and surfaces, disentangled from ambient lighting conditions. By stating the goal as such, the inverse graphics approach also makes firm commitments to the intermediate representations of the system (e.g., dividing the scene into layers; into surfaces and ambient lighting or atmosphere; etc.), the computational mechanisms required (e.g., Markov process to search the possibility space), and the benefits to the agent (e.g., lightness constancy; stable recognition under different lighting conditions, etc.). In contrast to traditional inverse graphics models, our framework replaces the high-level (cognitive, agentic) goal of discovering intrinsic object properties with a modular, encapsulated goal of recovering a complete representation of the image from the compressed code transmitted from the retina. Moreover, our implementation of edge-based reconstruction remains agnostic regarding the underlying intermediate representations and computational mechanisms — except that the required mechanisms must be implementable within a deep-convolutional / de-convolutional hierarchy. Empirically, we show that several effects often cited as evidence for scene factoring (e.g., Moon Illusion; Checkerboard Illusion) *can* emerge without explicit layered or lighting representations, suggesting that edge reconstruction can emulate the outcomes of such complex scene analysis.

It seems plausible that the mechanisms employed by the Edge-Net more closely resemble Grossberg and Todorović filling-in mechanisms than an inverse-graphics model, but we cannot rule out the possibility that the Edge-Net is implicitly extracting rich, structured representations that guide the reconstruction process. That is, it is possible that some operations, like “explaining away” dark patches based on cast shadows, or separating images into layers, or detecting local cues to transparency, etc., are in some way happening in the operations of the Edge-Net model. If so, the most parsimonious reading is that edge-based reconstruction provides a proximate training objective that induces useful decompositions when needed, without committing the system to explicit factorized scene representations or costly global inference.

Other seminal research on lightness perception has focused on developing cognitive theories that describe rules which explain how the human visual system constructs lightness percepts (Adelson, 2000; Gilchrist et al., 1999). These theories focus on perceptual phenomenology — e.g., which surface appears “white” or “black” in a scene. For example, Gilchrist has proposed an anchoring theory, in which the highest luminance region is treated as white across a wide range of absolute intensities (Gilchrist et al., 1999; Gilchrist, 2006). Critically, these anchoring phenomena appear to respect perceptual organization and grouping rules, with additional weight given to regions within the same plane or perceptual group (Gilchrist, 1977; Kingdom, 2011; Agostini & Galmonte, 1999). These anchoring accounts successfully predict phenomena like the Koffka illusion (Zdravković et al., 2012) and White’s illusion (White, 1979; Kingdom, 2011) — precisely the cases where Edge-Net currently fails. However, unlike the Edge-Net models, they are not image computable, and it is not clear how they would be computed over LGN-like edge contrast responses. Bridging the gap between image-computable models and these phenomenology-based theories is thus an important avenue for future research.

### Missing mechanisms and limitations

While the model showed many emergent illusions, there were clear cases when the model failed to capture human perceptual phenomenology. These discrepancies specifically occurred when there were multiple competing cues for the brightness of a bounded region. In a way, these errors have in common a missing account of figure-ground segmentation, or perhaps of “border ownership” for the edge responses (Nakayama et al., 1995). Relatedly, anchoring theories of lightness perception (Gilchrist et al., 1999; Gilchrist, 2006) explicitly take grouping into account, and can successfully predict phenomena like the Koffka illusion (Zdravković, Economou, & Gilchrist, 2012) and White’s illusion (White, 1979; Kingdom, 2011). This fail case for the Edge-Net model suggests that a grouping-dependent, segment-wise normalization (i.e., “anchoring”) is a *missing ingredient* in the edge-based reconstruction framework. And, as such, it provides a clear target for hypothesizing what the mechanism is, which can be tested by incorporating those modifications into the Edge-Net.

For example, possible light-touch architectural or loss-level changes that might support reconstructions that respect grouping based phenomena are: (1) a “no-new-edges” prior, to discourage introducing spurious contrast boundaries (e.g., implemented as edge-aware total variation or graph-Laplacian smoothness); (2) a segmentation-aware decoder that applies segment-wise gain/offset (an anchoring-style normalization) to resolve competing local cues in favor of perceptual groups; (3) a limited long-range interaction module (e.g., dilated convolutions or bounded-radius attention) to implement the kind of grouping and border-ownership signals implicated in Koffka/White; and (4) a global–local coherence constraint that enforces cross-scale consistency (e.g., tying fine-scale surface fills to coarse-scale layout), in line with evidence that cutting-edge vision models benefit from globally coherent, context-sensitive integration across scales (Doshi et al., 2025). Note that these additions preserve the core “edges-in, surfaces-out” objective we are endorsing here, but could bring the Edge-Net more closely in line with human perceptual illusions where competing cues are resolved.

## Conclusion

We found that a single computational objective — to reconstruct natural images from pseudo-LGN edge responses — yields a visual system that produces systematic reconstruction errors which mimic many classic human lightness illusions, including several that have been argued to depend on complex scene decomposition and analysis. These results suggest that human perceptual experience can be strongly shaped by a simple, modular goal to combine edge responses and learned natural-image priors to reconstruct complete surface representations, and that this goal alone is sufficient to explain (much of) why the world looks the way it does to us. More broadly, this deep learning framework provides a bridge between research on computational vision, visual neuroscience, and visual perception, and paves the way towards a unified theory that treats laboratory phenomena (e.g., lightness illusions) not as the objective outputs of visual processing, but as by-products of efficient coding algorithms adopted by biological visual systems.

## METHODS

### Section 1: Model Training

In PyTorch, we implemented a multi-resolution convolutional network (MCNN), an optimized U-Net architecture from Wang et al. (2020). This model consisted of a custom 8-layer encoder, a convolutional bridge that expanded the encoder’s output to 2048 channels, and an 8-layer decoder module. The primary reconstructive goal was to output full grayscale images from a Laplacian-filtered (edge-based) input of the same image at 512 by 512 resolution. The encoder included 8 convolutional layers where the first layer used a 31×31 kernel to capture global context. Subsequent layers downsampled the input using stride-2 convolutions, progressively increasing the channel depth to 1024. Symmetrical to the encoder, the decoder had skip connections from the corresponding encoder layers. The model produced grayscale reconstructions at each of the eight decoder layers, ranging from 4 by 4 to 512 by 512, with the last layer using a Sigmoid activation to produce pixel values between 0 and 1. These models were trained end-to-end in a supervised fashion to minimize the total mean absolute error – the sum of reconstruction errors across all output scales.

We sourced 4 datasets (1k_ds1, 1k_ds1, 8k_ds1, 8k_ds2) of natural images from ImageNet’s training split and one validation set (8k_val) from ImageNet’s validation split. Datasets 1k_ds1 and 1k_ds2 each contained 1024 images, split into 75% for training and 25% for validation. Datasets 8k_ds1 and 8k_ds2 each contained 8000 images and shared a validation set of 2048 images (8k_val). To assess dataset independence, we used a hash-based overlap analysis. Overlap across the 1k-sized datasets was negligible (only 2 images were shared across both datasets). Across both the ImageNet training and validation splits, we observed minimal image repetition: dataset 8k_ds1 and 8k_ds2 had 7998 and 7997 unique images respectively, with 7 images in common. The shared validation set of 8k_val overlapped with only two images from each of 8k_ds1 and 8k_ds2. Finally, the overlap between the 1k-sized and 8k-sized datasets was at most 33 images (<3.5% of the 1k dataset and <0.5% of the 8k dataset). These results confirm that the datasets were largely independent. For edge-based models, we trained 8 models with different initialization seeds on each dataset.

We trained variants of the model architecture on one alternate reconstructive goal – denoising. This denoising goal was to reconstruct clean grayscale images after adding Gaussian noise with either standard deviation of 0.1 or 0.3. We trained 8 models with different initialization seeds on datasets 1k_ds1 and 8k_ds1 for each noise level. We add Gaussian noise on the fly and the target was the original grayscale images.

Other than the reconstructive goals and specific input preprocessing, all models used the same training parameters. During training, we used an AdamW optimizer set at learning rate = 0.001, β_1_ = 0.9, β^2^ = 0.999 and ε = 1e-7. We used a L1 loss summed across the models’ reconstructions at every scale. All models were trained for 128 epochs with a batch size of 64. For splitting datasets 1k_a and 1k_b into train/validation splits or adding any noise, we used a fixed starting seed of 1234. During the validation phase of all denoising models, the noise seeds were offset from the training seed. Any model trained on a 8k dataset used a FFCV loader and edge inputs were calculated on the fly.

### Section 2: Stimuli Creation for Models

Preprocessing of the stimuli largely varied based on the reconstructive goal of the model. All images, during training and testing, were first ensured to be converted to grayscale, resized to be 512 by 512, and standardized between 0 and 1 before any filtering step. For the edge-based models, all stimuli were passed through a custom Laplacian kernel of size 3 by 3. This filter approximates the behavior of a center-surround receptive field of a retinal ganglion cell or even a lateral geniculate neuron. During training for all types of reconstructing and denoising models, a random horizontal flip was applied. For training edge-reconstructive models specifically, gaussian noise was added to the edge input to prevent overfitting, as following Wang et al., 2020. For testing and training for gaussian denoising models, all stimuli were first scaled between -1 to 1 before a Gaussian noise pattern was added (the result output was clipped between -1 and 1).

The three primary illusions in this study were the Craik-O’Brien-Cornsweet illusion, Anderson-Winawer Moon Illusion, and the Adelson Checkerboard Illusion. The stimuli for the Craik-O’Brien-Cornsweet Illusion and Anderson-Winawer Moon Illusion were hand-created in Python to create precise masks for the regions of interest. For the Anderson-Winawer Illusion in particular, we adapted Matlab code from Martin Hebart (http://martin-hebart.de/webpages/code/stimuli.html). We chose the Anderson-Winawer pattern at random, ensuring that its average pixel value or brightness within the disc was close to 127 (out 255). The resolution of all images was 512 by 512, as determined by the model’s architecture. Masks for the region of interests were created during the process of creating the illusions themselves, by keeping track of the pixels that remain constant in both illusory regions. The Adelson Checkerboard Illusion was directly adapted from the original illusion (https://persci.mit.edu/gallery/checkershadow/). Participants and models (*only during test phase*) all viewed the primary Anderson-Winawer Moon Illusion stimuli. To probe for robustness, participants and models also viewed variations of the Anderson-Winawer Illusion (see Stimuli under Behavioral Experiments below). For the edge-based models, we also conducted generalizability tests by creating 500 new hazes with similar average pixel value as our original test image.

We supplemented the Craik-O’Brien-Cornsweet illusion,Anderson-Winawer Illusion, and Adelson Checkerboard Illusion with illusions made in Photoshop or using the stimupy Python package (Schmittwilken et al., 2023) which allowed for parametric control and access to built-in ROI masks. From Purves et al. (1999), we directly adapted the Cornsweet Cube Illusion from (https://purveslab.net/downloads-2/#coc), used Photoshop to adapt the 3D Cornsweet - Curved Display, and used stimupy to adapt the 2D Cornsweet displays used in behavioral experiments. From Murray (2020), we directly implemented the Snake Illusion, Simultaneous Contrast Illusion (articulated version) and Checkerboard Assimilation Illusions present as part of the stimupy package. For the Haze Illusion and Koffka Ring Illusions, we used the versions of the illusions from Murray 2020 as a basis, but hand coded a version in stimupy to more closely match the distribution of pixels in the versions of the illusions presented in (Adelson, 2000). Finally, for the White Illusion, because we evaluated the effect of parametrically adjusting its width of the illusory rectangles on the models’ reconstructions, we used Murray (2020)’s illusion as a basis and developed a controlled set of illusions in stimupy. For all illusions, we ensured masks marked all illusory surfaces.

### Section 3: Behavioral Experiments

We had two independent groups of 30 participants for the Uniform Match Task and Pattern Match Task. Both tasks were hosted on the Prolific platform for recruiting participants and collecting data. Our inclusion criteria were: 1) participants from the US 2) first language being English. All participants were compensated at $12.00/hr. Our exclusion criterion is detailed as followed. Both experiments had six experimental conditions. If any participant’s responses are outside three standard deviations in three out of the six experimental conditions, we excluded them. No participant matched this standard. We only excluded one participant for the Pattern Match Task who reported they had technical difficulties and was unable to finish the task entirely. Our final sample size was 30 participants for the Uniform Match Task and 29 participants for the Pattern Match Task. Harvard Institutional Review Board considered these tasks as having minimal to no risk.

Both tasks used the same exact stimuli but varied in how the participant responded. We took the Anderson-Winawer Illusion presented to the models and adjusted the Michelson Contrast within the moon’s boundary such that we had three levels of contrast at 40%, 51%, and 62%. The Michelson Contrast was updated by scaling the pixel distribution by two factors that adjusted the mean and standard deviation respectively. With 3 contrast levels and 2 backgrounds (either light or dark haze), there were a total of 6 stimuli used in both tasks. Each stimulus was presented to the participant in 4 orientations, at 90 degrees intervals, to account for participant noise.

The Uniform Match Task was directly inspired by Anderson & Winawer (2005)’s primary task. The 6 illusion stimuli were presented at 4 orientations in a random order for each participant. Each trial presented 2 main panels to the participant: the left panel had a static display of one of the illusion stimuli and the right panel had an image of an equivalent size with a uniform gray disc in the center and fixed white noise in the background. Participants used a slider to change the brightness of the uniform disc from black to white to match their perceived brightness of the moon in the illusion. The illusion strength by stimulus was taken to be the difference in pixel value selected by the participant for the moon presented in the dark haze (*lighter moon*) – the moon presented in the light haze (*darker moon*) averaged across all presented orientation.

The Pattern Match Task differed from the Uniform Match Task in only one critical way – the participant’s response. Like with the Uniform Match Task, each illusion stimulus was presented in 4 orientations to the participants in a random manner. Each trial consisted of two subpanels – one displaying the original illusion and now the second panel displaying an image of equivalent size with the subsection of the moon (with the pattern) on top of fixed white noise. Participants used a 2D slider to change the brightness and sharpness of the texture within the moon simultaneously. Moving right increased the factor adjusting the mean of the pixel distribution (made it brighter) while moving left did the opposite. Moving upwards increased the factor adjusting the standard deviation of the pixel distribution (made it sharper) while moving down did the opposite. Each trial started at the midpoint of the grid – the moon texture presented in the Anderson-Winawer Illusion at 51% Michelson Contrast. The grid also contained the objective match to the moon presented in the illusion, but the participant was not informed as such.

For all illusions used in behavioral tasks, the same masks were used to evaluate the performance of models and participants for direct comparison. Because the behavioral tasks were conducted online, we could not control for luminance across participants or between outputs from the models and participants. As such, we measured illusion strength for each stimulus as the difference in mean pixel value of the selections chosen the participant for the *light moon* – *dark moon*, averaged across all presented orientations, as the most consistent and interpretable proxy for perceived brightness across models and participants.

### Section 4: Statistical Analysis of Illusion Strength in Model Reconstructions

For all models, we defined illusion strength by comparing the mean pixel values within the two regions of interest (ROIs) that are identical in the original stimulus but perceived differently Pixel values across all outputs were scaled between 0 and 255. An illusion was considered recapitulated if the mean difference in pixel intensity between the illusory-light ROI and illusory-dark ROI, was significantly positive in direction of the perceptual effect.

### 4.1 Generalization Across Edge-based Models and Gaussian Noise Models

To test the robustness of observed illusions across reconstructive objectives, we computed one sample t-tests for each set of independent edge-based models (separated by training dataset) and gaussian denoising models (separated by gaussian noise level) to evaluate whether the illusion magnitude was significantly greater than 0. For each gaussian denoising model, we first passed the illusion 10 times with different noise patterns at the specific level that the model was trained on and computed illusory effect as the mean pixel value difference in the illusory-darker ROI and illusory-lighter ROI in the average reconstruction. For each edge-based model, first, we calculated the illusory effect as the mean pixel value difference in the illusory-darker ROI and illusory-lighter ROI in the reconstruction output. For each gaussian denoising model, we computed the same illusory effect except on the average reconstruction after passing the stimulus through the model 10 times with independent noise patterns at the noise level. These analyses allow us to evaluate whether the illusion can consistently emerge across model instances and whether this effect is specific to edge-based reconstruction instead of reconstruction more broadly.

## Footnotes

1 Your visual system cannot see the model’s failure in Fig. 5 because you see the illusion!

