## Supplemental Info for "A Unified Account of Lightness Illusions via Edge-Based Reconstruction of Natural Images"

Supplemental Information for “Edge-based natural image reconstruction provides a unified account of many lightness illusions”

Srijani Saha, Talia Konkle, and George A. Alvarez

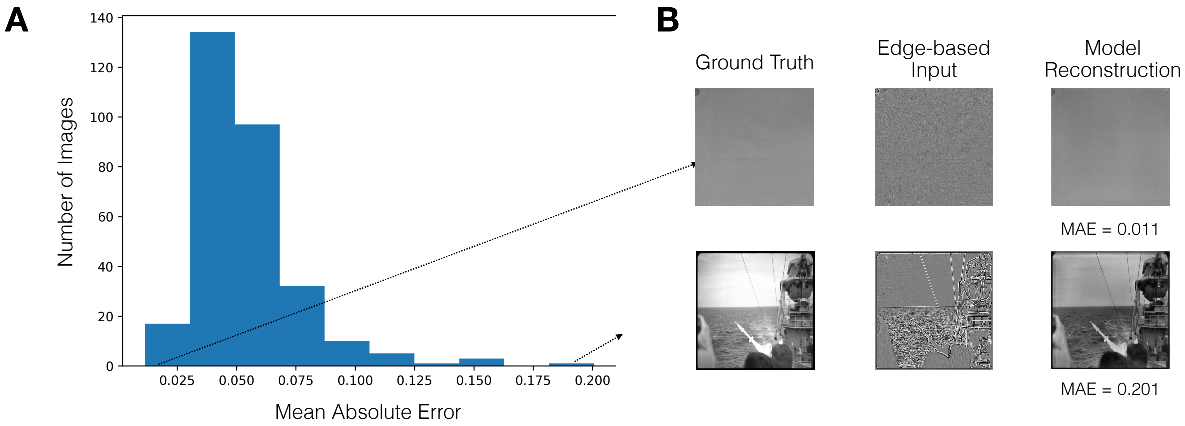


**Supplementary Figure 1: Reconstruction performance of the primary edge-based model**

Panel A plots the distribution of mean absolute error of the primary edge-based model’s reconstructions for an evaluation set of 300 new random ImageNet images from the ImageNet training subset. Panel B showcases the images with the best and worst reconstruction performance. We present the original image, the edge-based input fed into the model, and the model output or reconstruction. The model generally showed low reconstruction error across this set of natural images.

**Supplemental 2**: What are the reconstruction performances of the EdgeNets trained on 1k_ds1 and 1k_ds2?

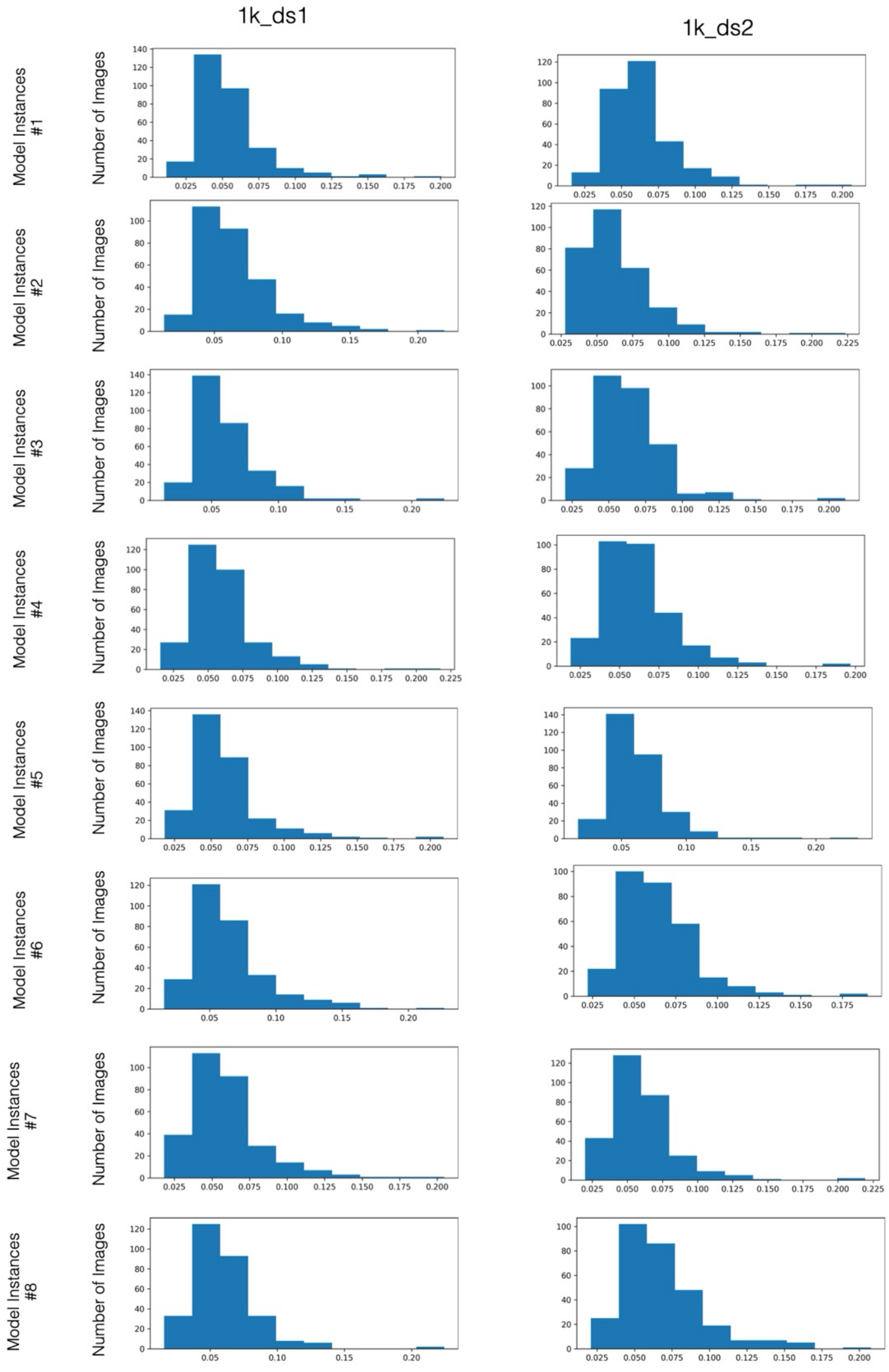

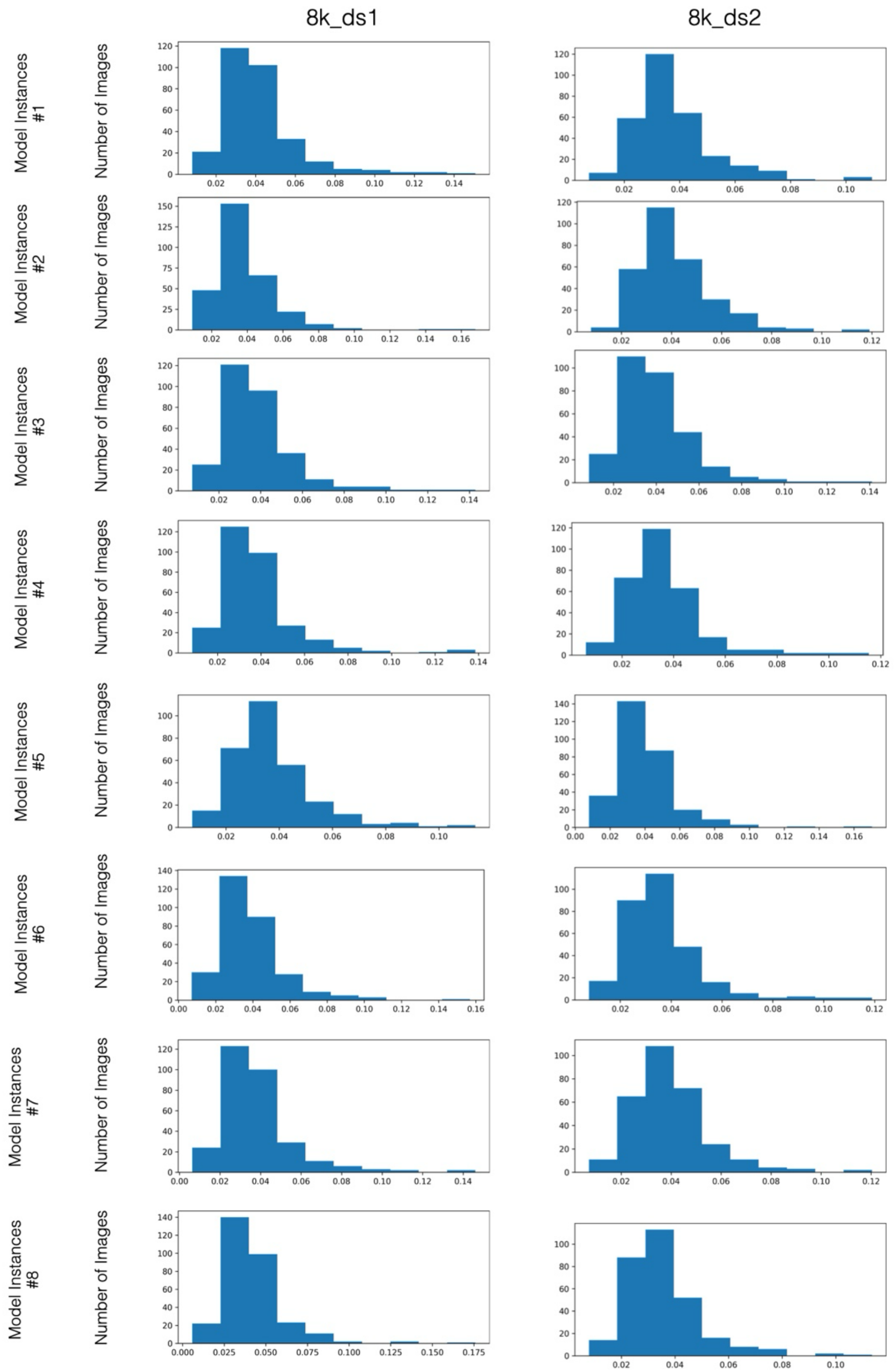


**Supplementary Figure 2: Distribution of reconstruction performance for Edge-based models**

Distributions of mean absolute error for the 32 edge-based models' reconstructions on the evaluation set of 300 new images. These distributions indicate consistent reconstruction performance across each set of edge-based models, qualitatively having better performance as the training set size increased.

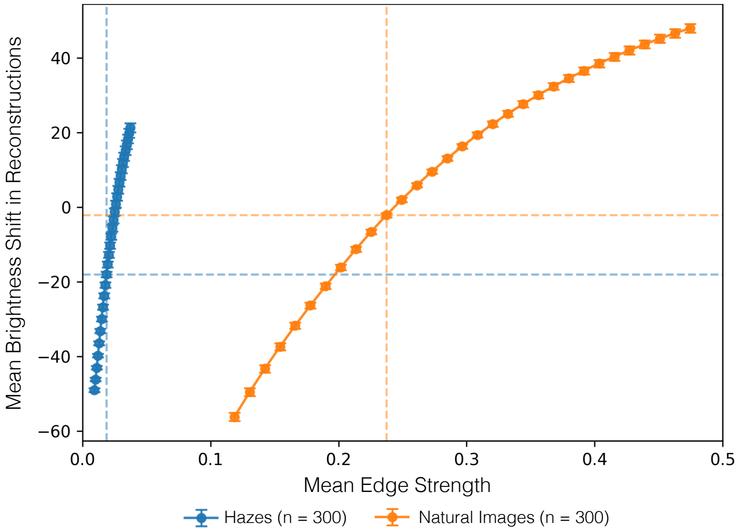


**Supplementary Figure 3: Effect of edge strength signal on the brightness in primary edge-based model's output**

The primary edge-based model's reconstructions of quasi-natural Adelson-Winawer haze images (n = 300) and natural ImageNet images (n = 300) became brighter as the mean edge strength of the input increased and became darker when the mean edge strength was weakened, which reflects the model's reliance on a darker prior when it has less edge information. Because hazes have structural differences from natural images, they do not overlap exactly.

Laplacian edge inputs of each set of hazes and natural images was uniformly scaled by a factor between 0.5 and 2.0. Across the images, the edge strength was calculated the average absolute Laplacian value and mean brightness shift was calculated as the average difference in mean pixel value between the reconstruction and the original image. Error bars represent the standard error across the 300 images and dashed lines mark the mean edge strength and mean brightness shift for the unscaled original inputs for the hazes and natural images.

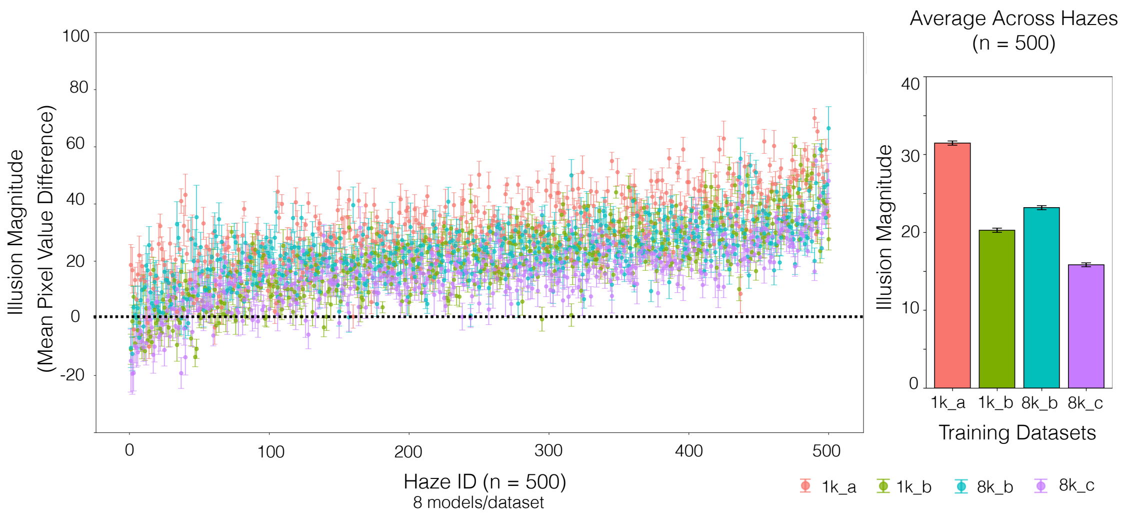


**Supplementary Figure 4: Robustness of Adelson-Winawer Illusion across edge-based models**

Mean illusion magnitude for 500 Anderson-Winawer Illusion Stimuli across 32 models (8 seeds x 4 training sets). The left panel plots the models' average illusion strength for each of the Anderson-Winawer Illusion stimulus, grouped by the 4 training datasets with error bars highlighting the variation across the 8 seeds. The right panel plots the average illusion strength across the 500 hazes for the 32 models, grouped by the 4 training datasets with error bars again representing variation across the 8 seeds. Although there is variation in illusion strength by individual haze structure, two-tailed sign-flip permutation tests for each set of models show that most models generally reproduced the illusion consistently (1k_a: 476/500; 1k_b: 383/500; 8k_b: 433/500; 8k_c: 377/500, $p$ <0.05).

trained on 8k_ds1 at different levels of Gaussian Noise?

**Distribution of Reconstruction Performance Across Test Set**
**(n = 300 ImageNet images)**

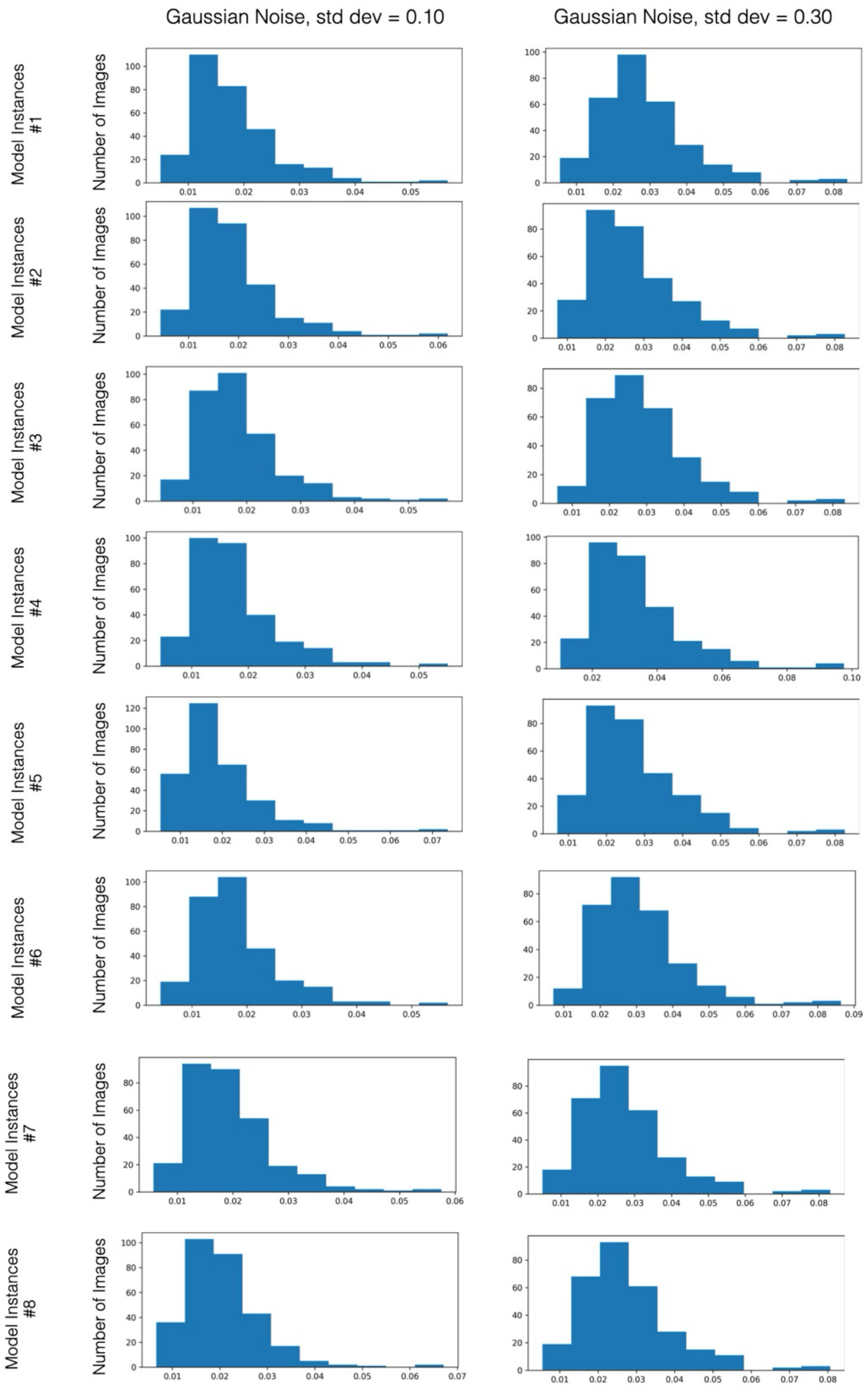

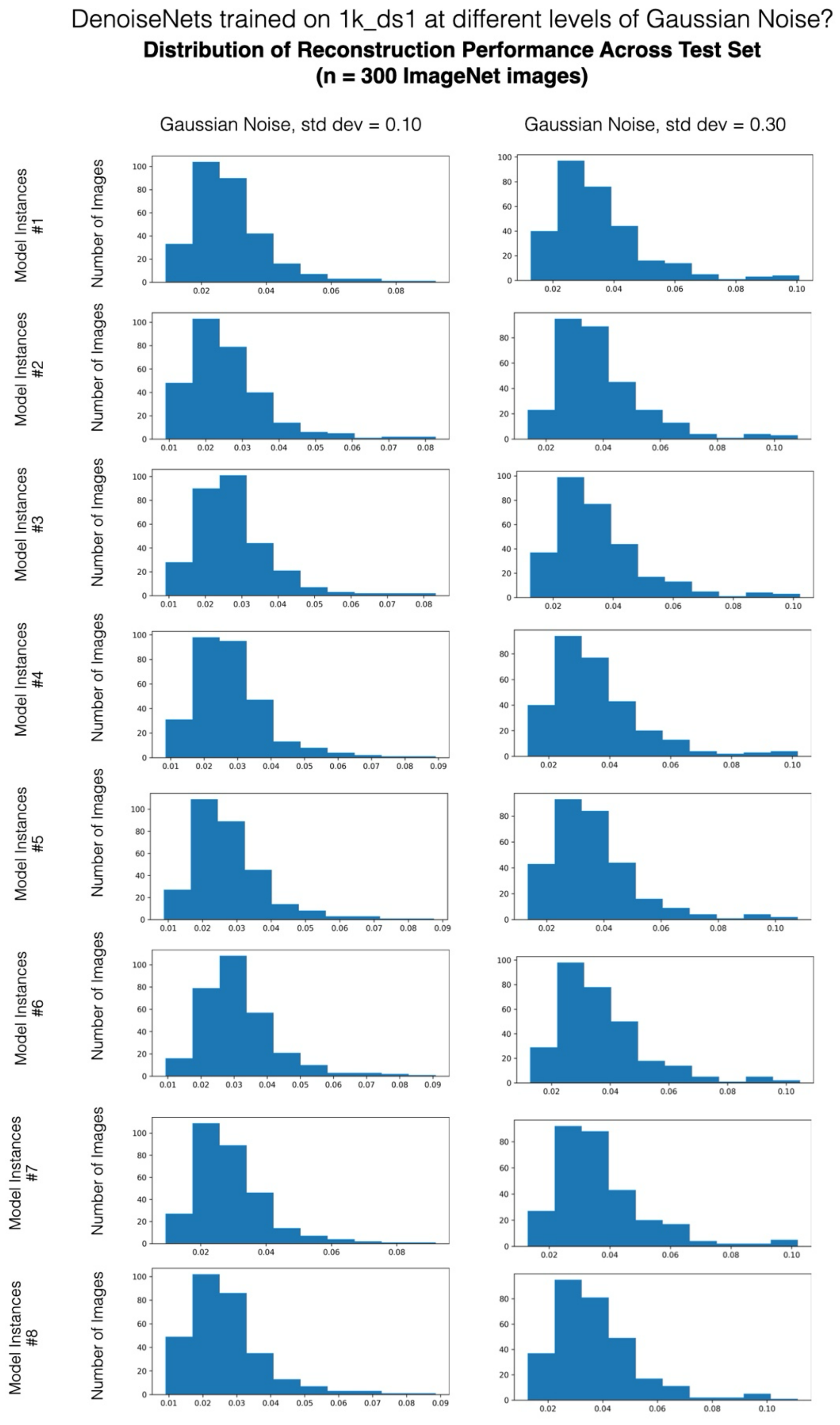


**Supplementary Figure 5: Distribution of reconstruction performance for Gaussian-denoising models (trained on 8k_ds1 and 1k_ds1)**

Distributions of mean absolute error for the 32 Gaussian-denoising models' reconstructions on the evaluation set of 300 new images. Each image was passed through the model 10 times with different gaussian noise patterns, and the mean absolute error was calculated the as the average reconstruction error across all the runs. These distributions indicate consistent reconstruction performance across the denoising models, qualitatively having better performance than the edge-based models trained on the smaller datasets and qualitatively similar performance to the edge-based models trained on larger datasets.

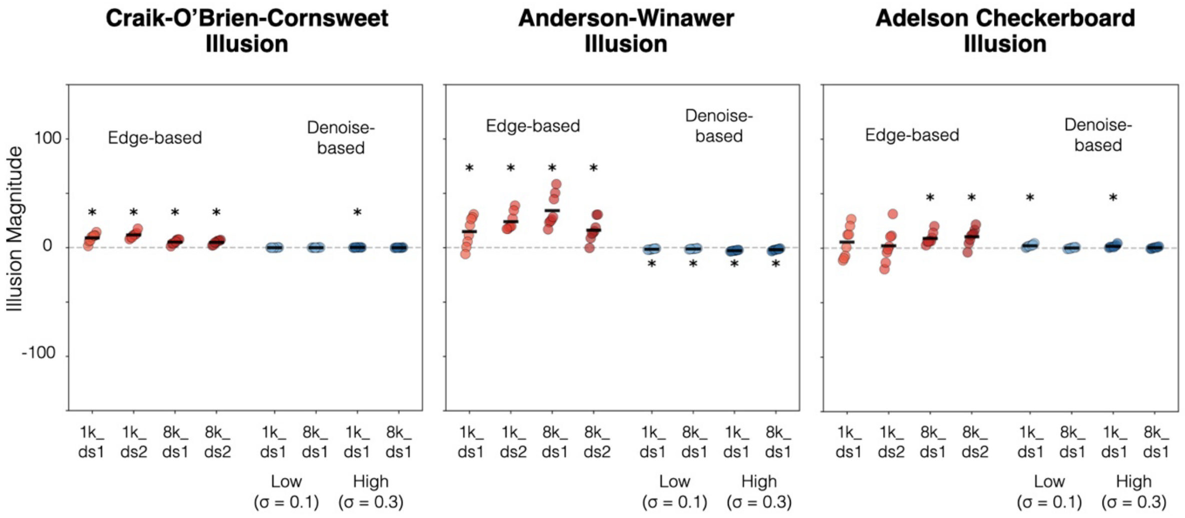


**Supplementary Figure 6: Comparison between edge-based models and Gaussian-denoising models for producing the core trio of illusions**

Figure 5 compares the mean illusion magnitude (plotted on the y-axis) between edge-based models (trained on 1k_ds1, 1k_ds2, 8k_ds1, and 8k_ds2) and Gaussian-denoising models (trained on 1k_ds1 and 8k_ds2) dataset for the core set of lightness illusions: Craik-O'Brien-Cornsweet Illusion, Anderson-Winawer Illusion, Adelson Checkerboard Illusion. Each dot represents the illusion magnitude for a particular model instance, and each bar represents a dataset mean. Across all illusions, Gaussian-denoising models did not exhibit an illusory effect consistently across datasets or level of Gaussian noise.

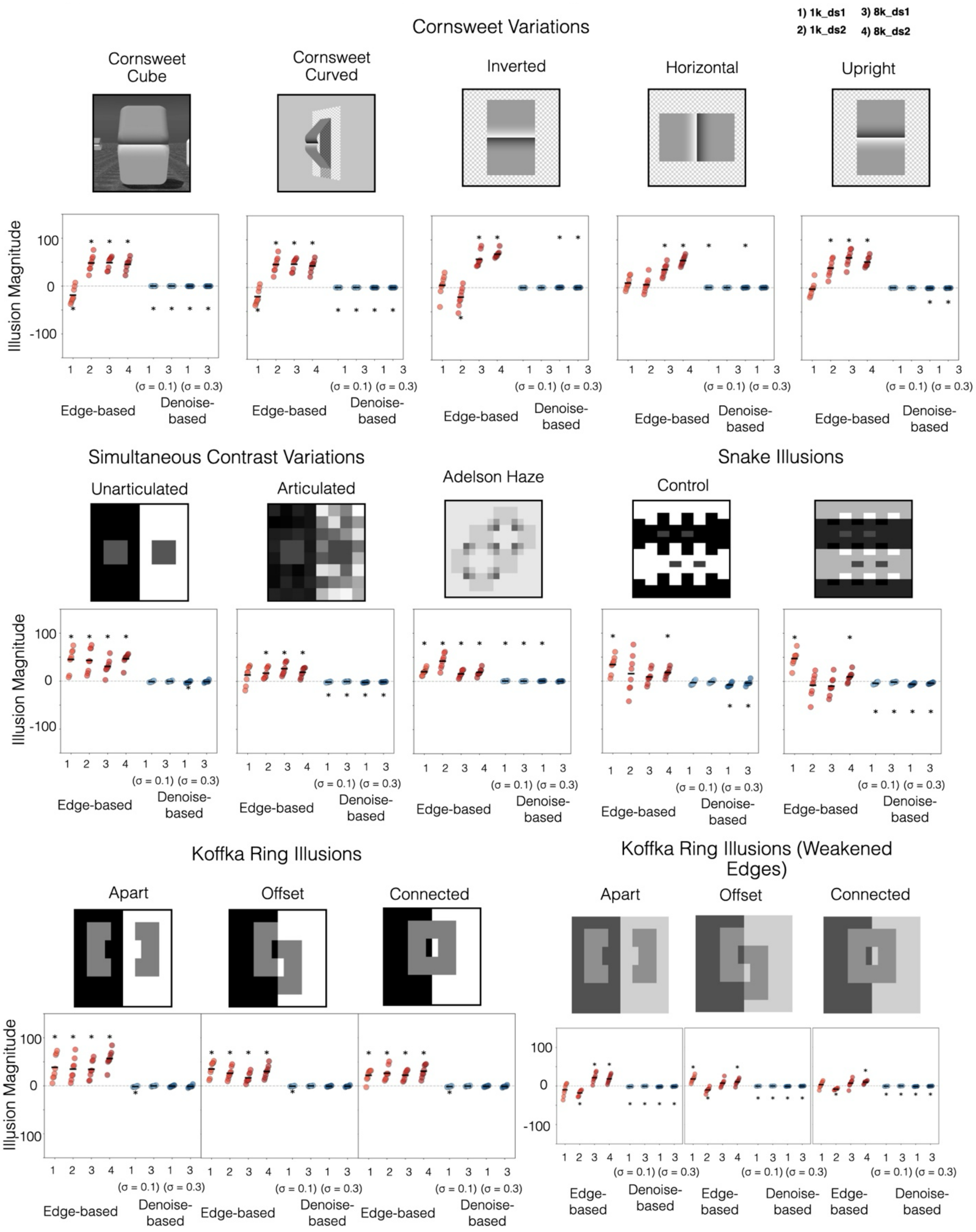
1) 1k_ds1
2) 1k_ds2
3) 8k_ds1
4) 8k_ds2
Cornsweet Variations
Cornsweet Cube
Cornsweet Curved
Inverted
Horizontal
Upright
Illusion Magnitude
100
0
-100
1 2 3 4 1 3 1 3
(σ = 0.1) (σ = 0.3)
Edge-based
Denoise-based
Simultaneous Contrast Variations
Unarticulated
Articulated
Adelson Haze
Snake Illusions
Control
Koffka Ring Illusions
Apart
Offset
Connected
Koffka Ring Illusions (Weakened Edges)

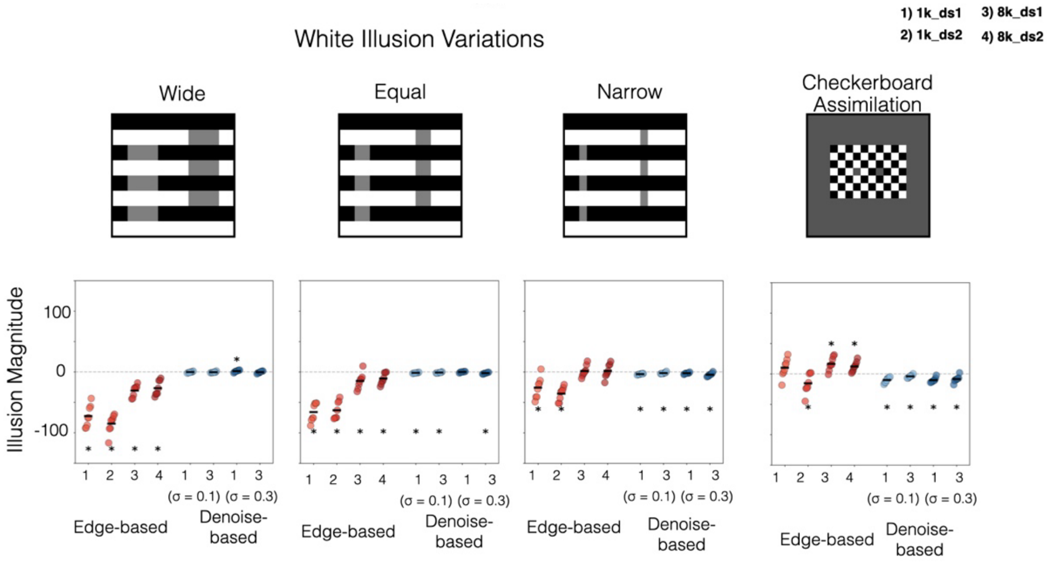


**Supplementary Figure 7: Comparison between edge-based models and Gaussian-denoising models for producing the additional set of lightness illusions**

Figure 6 compares the mean illusion magnitude (plotted on the y-axis) between edge-based models (trained on 1k_ds1, 1k_ds2, 8k_ds1, and 8k_ds2) and Gaussian-denoising models (trained on 1k_ds1 and 8k_ds2) dataset for the broader set of lightness illusions. Each dot represents the illusion magnitude for a particular model instance, and each bar represents a dataset mean. On the x axis, 1 stands in for 1k_s1, 2 stands in for 1k_ds2, and 3 stands in for 8k_ds1, and 4 stands in for 8k_ds4.  One sample t-tests on the illusion magnitude were used to evaluate if the mean illusion magnitude different significantly from zero. Broadly, the Gaussian-denoising models do not recapitulate lightness illusions across training datasets or level of Gaussian noise.

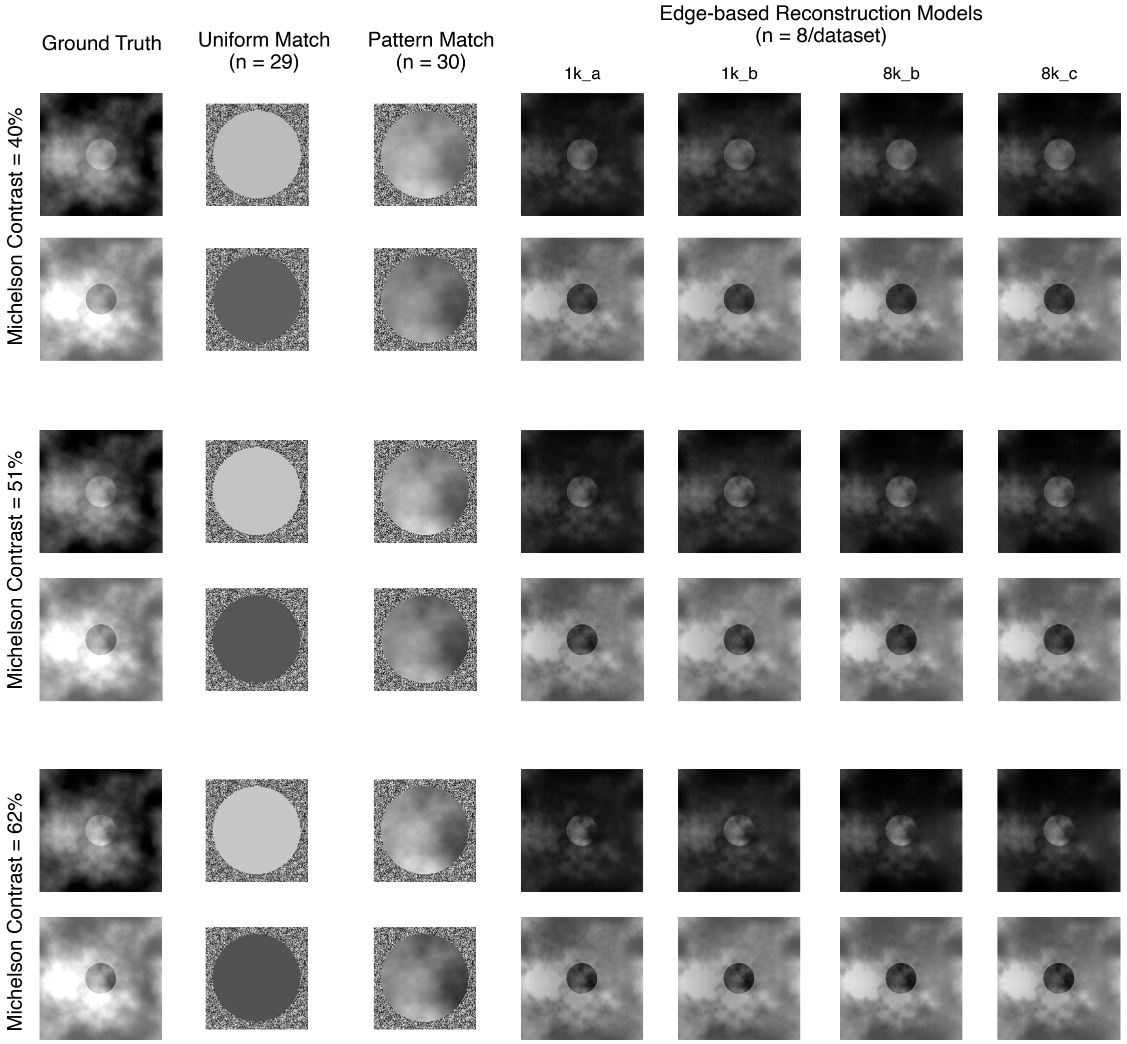


**Supplementary Figure 8: Visualization of the average edge-net model reconstructions and human participant responses for the Anderson-Winawer Illusion**

Average behavioral responses from human participants and reconstructions from edge-based models are shown for each level of Michelson Contrast of the Anderson-Winawer Illusion. Behavioral data is from the Uniform Match ($n$ = 29) and Pattern Match ($n$ = 30) tasks while the model reconstructions are the outputs averaged across the 8 models for 1k_ds1, 1k_ds2, 8k_ds1, and 8k_ds2. Irrespective of the behavioral task instructions or the dataset, participants and models perceive a brighter moon in the dark haze and a darker moon in the light haze.

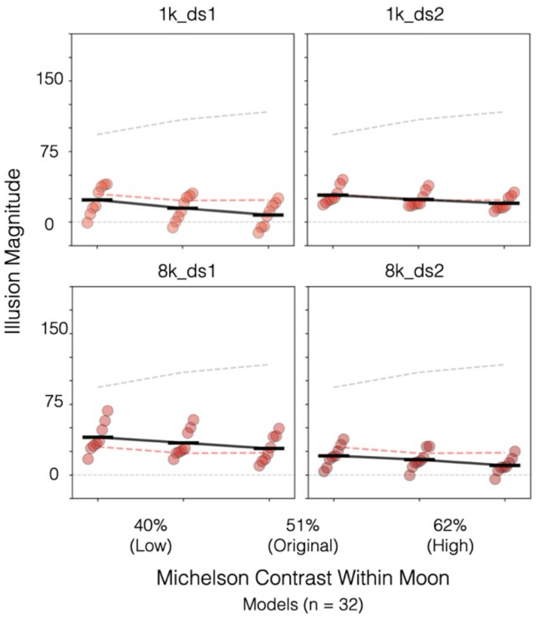


**Supplementary Figure 9: Human and Model Signatures of Anderson-Winawer Moon Illusion and the parametric effects of contrast across full set of models.** Each panel shows a comparison between average human behavior (uniform match: gray dashed line; pattern match: red dashed line) versus 8 edge-based models (red dots) for that dataset. The illusion magnitude (y-axis) is again plotted as a function of the Michelson Contrast (x-axis) of the moon in the target image for the same stimuli. Each dot represents a model instance, and each black bar represents the mean performance across models
